# Soil microbes prefer organic acids over sugars in simulated root exudation

**DOI:** 10.1101/2024.12.03.626624

**Authors:** Julia Wiesenbauer, Stefan Gorka, Kian Jenab, Raphael Schuster, Naresh Kumar, Cornelia Rottensteiner, Alexander König, Stephan Kraemer, Erich Inselsbacher, Christina Kaiser

## Abstract

Sugars and organic acids, primary components in plant root exudates, are thought to enhance microbial decomposition of organic matter in the rhizosphere. However, their specific impacts on microbial activity and nutrient mobilisation remain poorly understood. Here, we simulated passive root exudation to investigate the distinct effects of sugars and organic acids on microbial metabolism in the rhizosphere. We released ^13^C- labelled sugars and/or organic acids via reverse microdialysis into intact meadow and forest soils over 6-hours. We measured substrate-induced microbial respiration, soil organic matter mineralization, metabolite concentrations, and substrate incorporation into lipid-derived fatty acids. Our results reveal a pronounced microbial preference for organic acids over sugars, with organic acids being removed faster from the exudation spot and preferentially respired by microbes. Unlike sugars, organic acids increased concentrations of microbial metabolic byproducts and cations (K, Ca, Mg) near the exudation spot. Our results challenge the prevailing assumption that sugars are the most readily available and rapidly consumed substrates for soil microbes. Microbial preference for organic acids indicates a trade-off between rapid biomass growth and ATP yield. Our findings underscore the significant role of exudate composition in influencing microbial dynamics and nutrient availability, and emphasize the importance of biotic and abiotic feedback mechanisms in the rhizosphere in regulating root exudation.

## 1. Introduction

Plants release a considerable fraction of recently photo-assimilated carbon (C) as low-molecular weight compounds, such as sugars, organic acids and amino acids, via their fine roots (Badri and Vivanco, 2009; Jones et al., 2009; Canarini et al., 2019; Vives-Peris et al., 2020). These root exudates trigger a cascade of biotic and abiotic interactions in the rhizosphere. They facilitate the release of organic matter from mineral surfaces, and enhance its decomposition by soil microbes, thereby increasing nutrient availability in the soil immediately surrounding the roots (Brzostek et al., 2013; Drake et al., 2013; Kuzyakov et al., 2015; Mommer et al., 2016; Lu et al., 2019). Root exudation rates, which are linked to plant photosynthesis rates, appear to be increasing due to increasing atmospheric CO2 concentrations, with repercussions for soil organic matter turnover in the rhizosphere (Phillips et al., 2011; Meier et al., 2015; Calvo et al., 2017; Gargallo-Garriga et al., 2018; Dong et al., 2021).

Beyond just the quantity of root exudates, the chemical identity of exudates is expected to influence rhizosphere processes. Sugars and organic acids, two of the main compound classes in root exudates (Smith, 1976; Fender et al., 2013), are expected to affect the abiotic and biotic soil environment differently (Jones et al., 2003; Oburger et al., 2009; Steinauer et al., 2016). In contrast to sugars, organic acids are charged, and so they are more likely to be sorbed to the soil mineral phase, becoming unavailable to microbes until desorption (Jones and Edwards, 1998). Moreover, organic acids liberate organic compounds from mineral-organic associations, facilitating their subsequent decomposition by soil microbes (Keiluweit et al., 2015; Jilling et al., 2021). At the same time, they solubilise or desorb mineral-associated nutrients and metals through acidification, chelation, and exchange reactions (Keiluweit et al., 2015; Adeleke et al., 2017; Jilling et al., 2021). Sugars, on the other hand, are thought to be primarily and rapidly taken up and metabolised by microorganisms, with sorption, leaching and plant uptake playing a much smaller role (Gunina and Kuzyakov, 2015). When artificial root exudates were added to low fertility grassland soils, sugars increased respiration and dissolved organic C (DOC) more strongly than did organic acids (Liu et al., 2022). Additionally, an experiment applying different root exudate compounds to woodland soils showed that glucose was more strongly retained in microbial biomass than oxalic acid (Oldfield et al., 2018). Conversely, it is thought that organic acids cause stronger changes to the microbial community composition than sugars, as they are likely metabolised by specialised microorganisms, whereas sugars are utilised by a broader range of microbes (Landi et al., 2006; Eilers et al., 2010; Shi et al., 2011; Macias- Benitez et al., 2020).

Microbial growth on organic acids requires different metabolic pathways than those used when growing on sugars. Organic acids, such as short chain fatty acids, need to be transformed by specific enzymes into acetyl-coenzyme A, which is then fed into the Tricarboxylic Acid (TCA) cycle to obtain energy (Pavoncello et al., 2022). The production of precursors necessary for cellular biosynthesis requires: i) a ‘glyoxylate bypass’ in the TCA cycle to produce, e.g., amino acid precursors (Wolfe, 2005; Pavoncello et al., 2022) and ii) gluconeogenesis for the synthesis of glucose from non-carbohydrate precursors. The latter pathway runs in the opposite direction to glycolysis (breakdown of glucose to pyruvate) (Schink et al., 2022). Microbes may switch between sugar and organic acid metabolization, but the reversal of the central C flux direction, from glycolysis to gluconeogenesis, and vice versa, involves a lag phase of minutes to hours (Basan et al., 2020). On the one hand, a switch to gluconeogenesis can occur after the depletion of sugars that were initially metabolised by glycolysis. For example, *E. coli* excretes acetate during overflow metabolism as a strategy to maximise growth on glucose (Basan et al., 2015), and may later switch to utilizing this acetate once glucose is depleted (Wolfe, 2005). On the other hand, both metabolic pathways could run side by side, carried out by different parts of the microbial community specialised in the utilization of different substrates (cross-feeding), which would allow for a more resource-efficient utilization of available substrates within the microbial community. It is therefore unsurprising that substrate identity affects microbial community assembly (composition, diversity) and function (Shi et al., 2011; Steinauer et al., 2016; Zhalnina et al., 2018; Gu et al., 2020; Estrela et al., 2021).

To date, the magnitude and nature of the effects of root exudate composition on microbial activity is elusive. Little is known about the distinct effects of major compound classes, such as sugars or organic acids, on the dynamics of complex microbial communities in the rhizosphere. Although some studies have investigated the effect of different compound classes in artificial root exudates on rhizosphere processes, such as soil-mineral interactions (Keiluweit et al., 2015), soil organic C and nitrogen (N) turnover (Chari and Taylor, 2022; Liu et al., 2022) and greenhouse gas fluxes (Girkin et al., 2018a, 2018b), we still lack a deeper understanding of how rhizosphere communities utilise different compound classes within exudates.

Studying microbial processes in undisturbed soil is crucial for obtaining accurate and ecologically relevant insights, as biogeochemical and structural heterogeneities at the pore-scale govern soil processes and functions (Schlüter et al., 2020). Intact soil structure preserves the natural spatial distribution of resources and decomposers, significantly influencing microbial access to resources and thereby affecting microbial activity and decomposition processes (Nunan et al., 2020). Equally important is the accurate simulation of passive root exudation. Artificial ‘root exudates’ can be introduced into the soil using various techniques beyond directly mixing substrates into the soil. Methods include the addition of ‘exudates’ with pipettes or needles (Luo et al., 2014; Steinauer et al., 2016; Liu et al., 2022), delivery through microporous capillaries (Baumert et al., 2018; Sokol and Bradford, 2019; X.

Zhang et al., 2019; Chari and Taylor, 2022) or by an automated drip-tip system (ARES) (Lopez-Sangil et al., 2017). These so-called ‘artificial roots’ often combine exudate release with water flow. However, most root exudates are passively released (Canarini et al., 2019), thereby depending on diffusion gradients, which create a direct feedback between their removal by biotic and abiotic soil processes and exudate release rates. Conventional mass flow-based ‘artificial roots’ cannot account for this feedback, highlighting the need for methods that better simulate natural passive exudation.

By contrast, ‘reverse microdialysis’, which relies on passive diffusion through a small permeable membrane, enables the release of compounds from the microdialysis probe based on concentration gradients between the membrane’s interior and the surrounding soil solution (Buckley et al., 2022; König et al., 2022; Wiesenbauer et al., 2024). This method provides a more realistic simulation of passive root exudation by modelling how rhizosphere processes affect compound release rates. Specifically, if these compounds are not removed from the soil solution by biotic or abiotic processes, their accumulation inhibits further diffusion, highlighting the dynamic feedback from the rhizosphere. Measuring this ‘uptake’ feedback not only offers insights into processes that selectively remove certain compounds from the rhizosphere but may also inform about the capacity for passive exudate release by plant roots under specific conditions.

Our previous work showed that organic acids were released more rapidly than sugars in soil, suggesting quicker removal from the rhizosphere compared to sugars (König et al., 2022; Wiesenbauer et al., 2024). We showed that this faster removal cannot be attributed to molecular size, which primarily governs diffusion in water. Instead, it may result from a more rapid adsorption to soil minerals or a microbial preference for metabolizing organic acids over sugars. This finding contradicts the general consensus that simple sugars are more rapidly metabolised by soil microbes due to their high energy yield (Gunina and Kuzyakov, 2015; Schink et al., 2022). To determine whether abiotic processes or microbial preferences for metabolising organic acids drive these differences in the rhizosphere, it is necessary to measure microbial utilisation of different compound classes individually. This approach would enhance our understanding of how different compound classes within root exudates contribute to nutrient mobilisation and C turnover in the rhizosphere.

The aim of this study was to analyse the distinct effects of sugars and organic acids in artificial root exudates on fine-scale temporal dynamics of microbial activity and soil chemistry at root exudation hotspots in intact soil. Specifically, we aimed to differentiate the impact of these compound classes on microbial and abiotic rhizosphere processes at root exudation hotspots using two distinctly different soil types — beech forest soil and extensively managed meadow soil — allowing for a broader perspective. To accomplish this, we released ^13^C-labelled sugars and organic acids, both individually and in combination, into intact soil cores by reverse microdialysis over a 6-hour period while simultaneously collecting metabolites from the soil solution for 18 days. We measured substrate-induced respiration and soil organic matter (SOM) mineralization, and the incorporation of substrates into microbial biomass (lipid-derived fatty acids). Our results show that soil microbes preferentially utilise organic acids over sugars in root exudates. This suggests that organic acids are vital not only for mobilizing organic compounds from soil minerals but also serve as a primary C and energy source for rhizosphere microbes, potentially even more important than simple sugars.

## 2. Materials and Methods

### 2.1 Soil sampling

Soil samples were collected from an extensively managed meadow (Gumpenstein, Austria, 47° 29’ N, 14° 06’ E, 732 m asl.) on 11^th^ July, 2019, which was a Cambisol with loamy sand texture (Reinthaler et al., 2021), and a mature beech (*Fagus sylvatica*) forest (Klausen-Leopoldsdorf, Austria, 48° 07′ N, 16° 03′ E, 510 m asl.) on 23^rd^ July, 2019, which was a Dystric Cambisol with silty clay texture (König et al., 2022) (Tab. 1). For both sites, the upper 6 cm of soil was sampled after removing the above-ground biomass and the litter layer. Samples were taken at five spots along an 8 m transect in the meadow soil and five random spots within a 5 x 5 m plot in the beech forest, using large soil cores (10 cm diameter, 6 cm height).

**Table 1.**
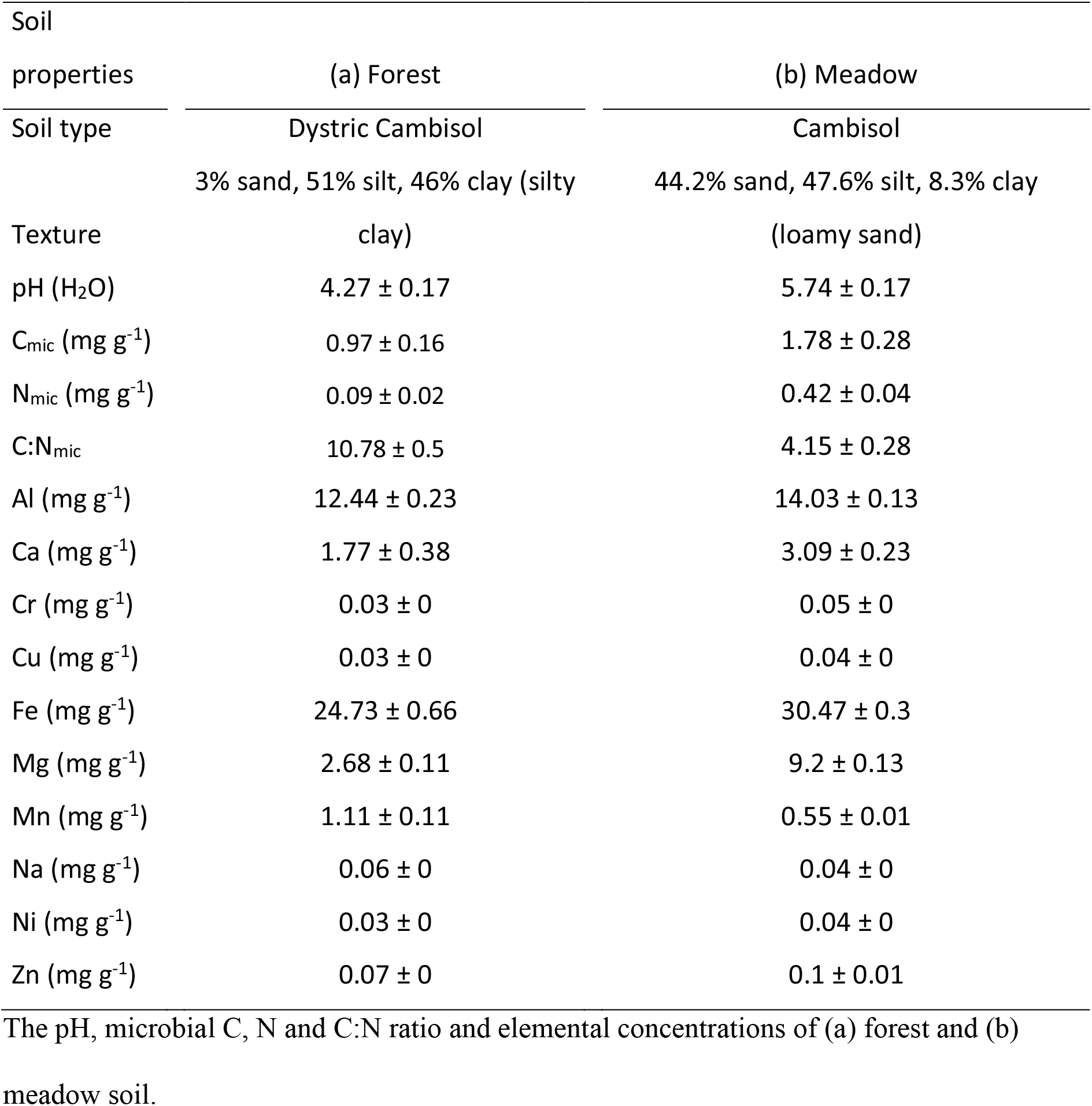
Soil properties (mean ± SE) of forest and meadow soils.

Initial soil analyses were conducted on samples sieved to 2 mm, collected adjacent to the soil cores used for the microdialysis experiment. These analyses included measurements of pH in soil slurries (water), and microbial C and N by chloroform fumigation extraction (Tab. 1, Methods S1). Total element concentration (Al, Ca, Cr, Cu, Fe, Mg, Mn, Na, Ni, Zn) of acid-digested (Aqua regia), finely ground, lyophilised soils was measured by inductively coupled plasma optical emission spectroscopy (ICP-OES, 5110, Agilent Technologies, USA) (Tab. 1).

We determined gravimetric water content of soils sampled with a small soil corer (1 cm diameter, 3 cm height) from larger cores. After removing roots, stones and organic material with tweezers, the soil was dried overnight in a drying oven. Initial gravimetric water content was 19 ± 3% (mean ± SD) in meadow and 12 ± 3% in forest soils. Water content of the large soil cores was adjusted to 30% during the 2-3 days post-sampling while the soils were incubated at field temperature (12 °C; until experiment).

### 2.2 Experimental design

To investigate the response of rhizosphere processes to simulated root exudate input in intact soil, we designed a microdialysis experiment using the large soil cores that preserved the natural soil structure. Three days after collection, four pairs of smaller soil cores were cored from each large soil core with stainless-steel cylindrical corers (1 cm diameter, 3 cm height). Each pair of cores – twin cores – was used for one of four experimental treatments.

These twin cores were treated as a single analytical unit, placed in a customised setup, termed a ‘mesocosm’, to ensure unified measurements of respiration, collection of analytes (‘dialysates’) and soil harvesting.

The twin-core approach was a modification from our previous study which used a single large core (2.8 x 3 cm) (König et al., 2022; Wiesenbauer et al., 2024). By using smaller soil cores, we effectively reduced the background respiration of soil outside the microdialysis sphere of influence, enhancing the sensitivity and accuracy of our respiration measurements.

For the mesocosm assembly, we used 50 ml Falcon tubes, modified by turning them on their lid and cutting them off at 4 cm height, with a Styrofoam placed into the lid to securely hold the twin soil cores in their stainless-steel corers (Fig. S1). Into the centre of each small soil core, a microdialysis probe (CMA 20, CMA Microdialysis AB, Kista, Sweden) with a membrane length of 10 mm and a molecular weight cut-off of 20 kDa was inserted.

The microdialysis probes were connected to syringe pumps (CMA 4004, CMA Microdialysis AB, Solna, Sweden), set to perfuse at a rate of 2.5 µl min^-1^ per membrane (the liquid is called perfusate). Additionally, they were connected to cooled (6 °C) fraction collectors (CM4 470, CMA Microdialysis AB, Solna, Sweden), which collected dialysate samples (i.e., the liquid containing the analytes collected through passive diffusion from the soil solution) in fractions of 45 min (112.5 µl). The setup allowed for the combined sampling of dialysates from the twin cores, ensuring a comprehensive assessment of nutrient dynamics within the mesocosm.

### 2.3 Simulation of root exudation using microdialysis

On day 1 of our 18-day long microdialysis experiment, we simulated a 6-hour long root exudation using the ‘reverse’ microdialysis approach (König et al., 2022). To investigate the effect of different root exudate compositions on soil metabolite dynamics, we utilised three artificial exudates (i.e., perfusates): pure sugar, pure organic acids, a combination of both, alongside a water-only control for comparison. Specifically, sugar-only exudates consisted of glucose and fructose, organic acid-only exudates contained acetic and succinic acid, and the mixed exudate was comprised of all four compounds, each uniformly labelled with 99 atom% ^13^C (Sigma-Aldrich). A C concentration of 500 µmol C l^-1^ per compound was maintained in each exudate treatment to facilitate direct comparisons of individual compounds across different exudate compositions. These compounds, commonly found in root exudates (Smith, 1976), were selected to cover a range of chemical structures typical of root exudates (Vives-Peris et al., 2020).

Initially, all mesocosms were infused with ultrapure water for 3 hours to assess the initial state of soil solution chemistry. Subsequently, each mesocosm was infused with its respective perfusate, i.e., either artificial root exudate (sugars, organic acids, mixed) or ultrapure water for controls. Following the 6-hour exudate simulation, all treatments were continuously infused with ultrapure water for three days, during which dialysates were collected to further explore soil metabolite dynamics. All dialysates were pooled in 3-hour fractions (four consecutive 45-minute dialysates) and stored at -20 °C until analysis. After this period, the pumps were turned off, and the membranes were left in the soils.

Subsequently, on days 4, 5, 6 and 18, the pumps were turned on for 4.5 hours each day to collect additional dialysates.

### 2.4 Soil metabolite analysis

Concentrations of sugars, organic and inorganic anions, and cations in the dialysates were quantified using high-performance liquid chromatography (HPLC, Dionex ICS 5000+, Thermo Fisher, Germany). Sugars (supplements: glucose, fructose; not detectable: galactose, sucrose) were measured on a Thermo CarboPac PA20 (0.4 x 150 mm) column with a Thermo CarboPac PA20G (0.4 x 35 mm) guard column at a constant flow rate of 8 µl min^-1^ with a KOH solvent. Anions (butyrate, lactate, propionate; supplements: acetate, succinate; no significant response: citrate, formate, malate, nitrate, oxalate, phosphate, sulfate) were measured on a Dionex IonPac AS11-HC (2 x 250 mm) column with a Dionex IonPac AG11- HC (2 x 50 mm) guard column at a constant flow rate of 0.25 ml min^-1^ with a KOH solvent. Cations (ammonium, potassium, magnesium, calcium; not quantifiable: manganese) were measured on a Dionex IonPac CS16 (5 x 250 mm) column with a guard column at a constant flow rate of 1 ml min^-1^ with a methansulfonic acid as solvent. Further details on the HPLC methods are provided in supporting information (Methods S2, S3, S4). Sugars and anions were measured at 14 time points, while cations were measured at 8 time points throughout the experiment.

### 2.5 Transfer and retrieval rates

We determined ‘transfer rates’ of glucose, fructose, acetate and succinate into the soil as the percentage of the total concentration of each compound in the exudation solution that was passively released into the soil. Transfer rates of compounds into water are primarily governed by diffusive molecular properties like size and charge, whereas in intact soils, transfer rates are significantly lower and predominantly determined by soil type, independent of molecular properties (König et al, 2022). This is because the soil’s physical structure confines the diffusible space for released molecules to a small volume around the membrane. Molecules that are not removed by biotic or abiotic mechanisms quickly accumulate outside the membrane, restricting further diffusion. Determining passive transfer rates of certain molecules into a given soil will thus inform about how readily these compounds are removed from the surrounding of the exudation spot by abiotic or biotic mechanisms. We calculated passive transfer rates based on the compound concentrations in the exudation solution before and after their passage through the microdialysis system (König et al., 2022).

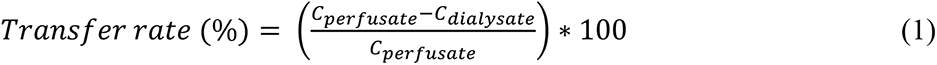

where C_perfusate_ is the concentration in the exudation solution before it was pumped through the microdialysis system (‘perfusate’) and Cdialysate is the concentration in the exudation solution that was collected after it was pumped through the microdialysis system (‘dialysate’).

Additionally, retrieval rates were calculated as the percentage of released compounds that were recovered from the soil surrounding the membrane over a 12-hour period following the exudation. For this time period, all treatments were infused with ultra-pure water to collect compounds present in the soil solution around the membrane.

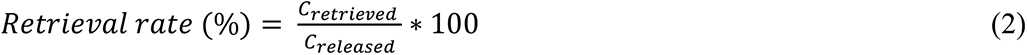

Here, C_released_ is the total amount of a compound released during the 6-hour exudation pulse and C_retrieved_ is the amount recovered from soil within 12 hours. Similarly to transfer rates, retrieval rates also provide information on the removal dynamics of released compounds. The lower the retrieval rates, the more of the released substances have already been removed from the spot by abiotic or biotic processes.

### 2.6 Respiration measurements

To measure microbial respiration, mesocosms were placed in airtight jars fitted with septa for gas sampling (Fig. S1). Microdialysis tubing was passed through the septa to enable continuous dialysate collection. Gas samples were taken on day 1 before, during, and after the substrate pulse, with additional samples collected on days 2, 4, 6, and 18. For gas sampling, the jars were closed, the first gas sample was taken with a syringe (18 ml) and replaced with artificial air (200 ppm CO2) to prevent negative atmospheric pressure. After incubation (∼90 min on day 1 & 2; during exudation ∼180 min for forest and ∼130 min for meadow), a second gas sample was taken, and jars were opened. On days 4, 6, and 18, incubation lasted ∼140 min, except ∼180 min for meadow soils on day 8. We measured CO2 concentration and ^13^C signature of gas samples using a headspace gas sampler (GasBench II, Thermo Fisher Scientific, Bremen, Germany) coupled to an isotope ratio mass spectrometer (Delta V Advantage, Thermo Electron, Bremen, Germany). The CO2 concentration and ^13^C signature of the second gas sample were corrected for added artificial air.

The respiration rate and its atom percent ^13^C (at% ^13^C) signature were calculated as the difference of ^12^C-CO2 and ^13^C-CO2 concentration between the first and second corrected gas sample. Using uniformly ^13^C-labelled substrates allowed us to distinguish between substrate-derived and SOM-derived respiration using a two-pool mixing model, considering the ^13^C signature of added substrate and natural ^13^C abundance of control mesocosm respiration. Detailed calculations are provided in the supporting information (Methods S5).

### 2.7 Harvest

On day 18, we harvested the soil surrounding the microdialysis membrane from two distances: ’central’ (≤ 2.5 mm radius) and ‘surrounding’ soil (> 2.5 mm radius). After pulling out the microdialysis probe, we used a 5 mm inner diameter stainless steel tube to collect the ‘central’ soil. Soil of twin cores were pooled for each mesocosm, and small stones and larger organic materials were removed using tweezers. Soils were lyophilised for 48 hours and stored at -20°C until further analysis. The ‘surrounding’ soil (> 2.5 mm from input) was used for lipid extraction, because the ‘central’ soil was used up in prior experimental work.

Consequently, the relatively low ^13^C signal observed is a conservative estimate of lipid-derived fatty acid enrichment near the membrane, as we previously demonstrated that ^13^C enrichment decreases significantly with distance from the microdialysis membrane (Wiesenbauer et al., 2024).

### 2.8 PLFA and NLFA analyses

To assess the incorporation of labile substrates into microbial biomass and their impact on microbial community structure, we extracted the phospholipid and neutral lipid fatty acids (PLFA, NLFA) from the lyophilised ’surrounding’ soils (> 2.5 mm from input) as described in Gorka *et al*. (2023). For a complete description of the extraction procedure and fatty acid assignments see the supporting information (Methods S6). The extracted fatty acid methyl esters were analysed using gas chromatography (Trace GC Ultra, Thermo Scientific, Germany) coupled to a mass spectrometer (ISQ, Thermo Scientific, Germany) for fatty acid identification and quantification, as well as a GC-Ultra (Thermo Fisher Scientific, Milan, Italy) coupled to an isotope ratio mass spectrometer (IRMS; Finnigan Delta-V, Thermo Fisher Scientific, Bremen, Germany) for determination of isotopic ^13^C/^12^C ratios.

### 2.9 Statistical analysis

All statistical analyses were performed using R version 4.2.2 (R Core Team, 2022). To determine significant (α = 0.05) differences between treatments, we conducted Kruskal-Wallis tests followed by Dunn’s tests as post-hoc tests (Bonferroni corrected p-values). We used this approach to assess differences between treatments for each time point in SOM-derived and substrate-derived respiration rates, compound concentrations in dialysate, transfer and retrieval rates, and PLFAs and NLFAs (nmol C g^-1^ DW, at% ^13^C). Figures were created using ggplot2 (Wickham, 2016).

## 3. Results

### 3.1 Microbes preferentially respired organic acids compared to sugars

Both forest and meadow soils exhibited higher respiration rates of organic acids compared to sugars (Fig. 1). Substrate addition did not significantly change SOM-derived respiration in either soil with SOM-derived respiration rates remaining similar to controls throughout the 18-day experiment (Fig. 1, Tab. S1). Both soils exhibited the highest substrate-derived respiration rate during the exudation pulse, which decreased thereafter (Fig. 1). Substrate-derived respiration rates were about 4-8 times higher when organic acids were part of the exudate solution compared to when only sugars were present. There was a minor, non-significant difference in substrate-derived respiration rates between soils receiving sugars and organic acids combined and those receiving only organic acids, which was slightly more pronounced in the meadow soil (Fig. 1, Tab. S1). Substrate respiration was measured for a 3-hour period during the 6-hour input pulse. Extrapolating the measured values to 6 hours, and relating them to the measured C input, shows that about 8, 10 and 9% of the released C has been respired during the pulse in the sugar, organic acid and mixed treatment, respectively.

**Figure 1.**
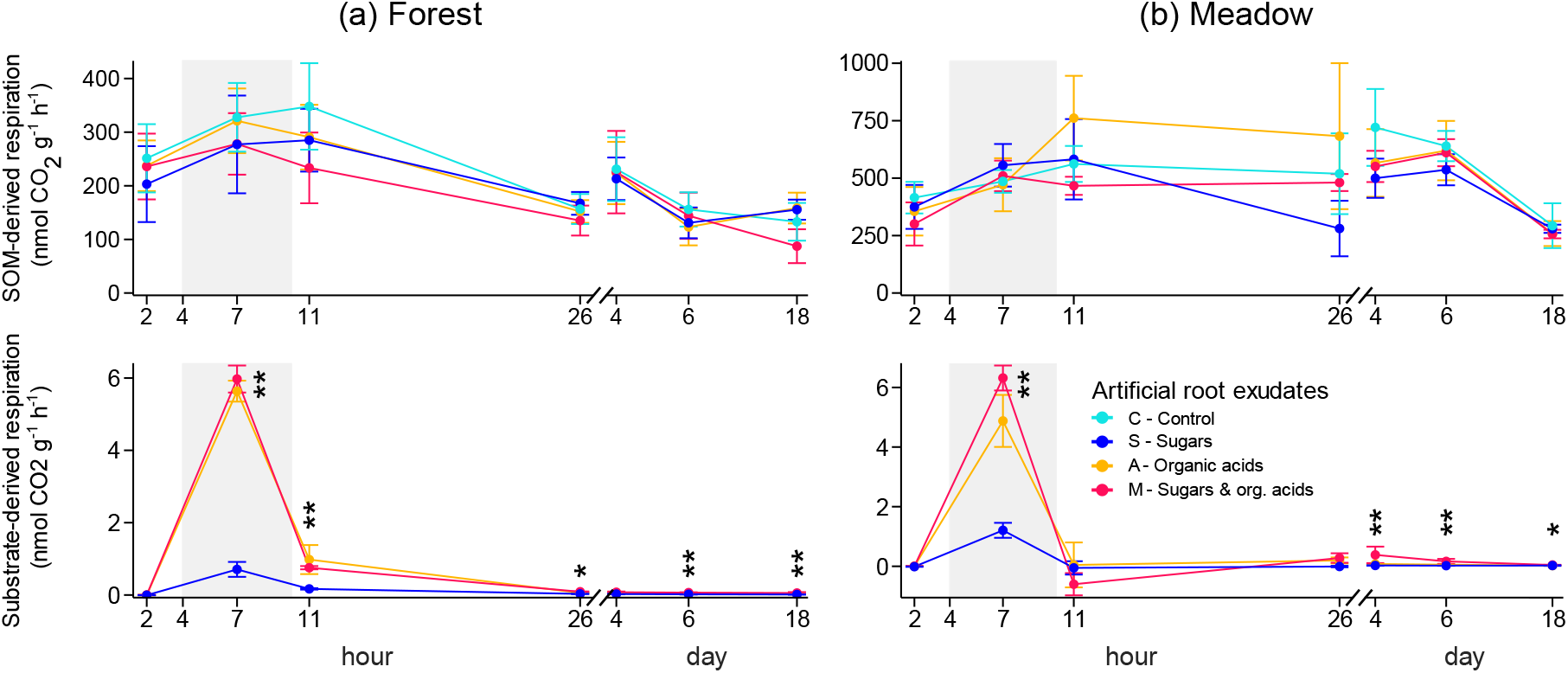
The SOM-derived and substrate-derived respiration rates (nmol CO2 g^-1^ dw h^-1^) for forest and (**b**) meadow soils. The 6-hour long labile substrate pulse (hours 4-9) is indicated by a grey background. An x-axis break follows the initial 26-hour period, and subsequent measurements (days 4-18) are presented on a day-timescale. Data are presented as means ± SE (*n* = 5). Asterisks denote significant differences (Kruskal-Wallis test, * < 0.05, ** < 0.01, *** < 0.001) between soils that received an input of sugars (dark blue), organic acids (orange), sugars combined with organic acids (pink), and the control (light blue; only SOM-derived respiration). Post-hoc test results of substrate-derived respiration are provided in Table S1.

### 3.2 Higher passive release for organic acids than sugars

We observed that the transfer rates (percentage of artificial root exudates, i.e., perfusates, inside the microdialysis membrane that were passively released, i.e., diffused, into the soil) were considerably higher for organic acids (55-67% acetate, 39-41% succinate) than for sugars (7-17% glucose, 14-23% fructose) (Fig. 2). Additionally, the majority of the transfer rates were higher in the first half compared to the second half of the 6-hour long labile substrate pulse (Fig. S2). Given that each compound was present at identical concentration (500 µmol C^-1^ l^-1^ per compound) in the perfusates, differences in transfer rates reflected differences in absolute amounts of C released. Over the 6-hour input period, on average 90-110 nmol C was released from the sugar-only perfusate, while 430-490 nmol C was released from the organic acids-only perfusate (Fig. 3). The mixed sugar and organic-acid treatment released about 550-660 nmol C, reflecting the combined sum of the C released in the sugar-only and organic acids-only treatments. There was no significant difference in the amount of each compound released, regardless of whether they were in a single-compound class treatment or the mixed treatment (Fig. 3).

**Figure 2.**
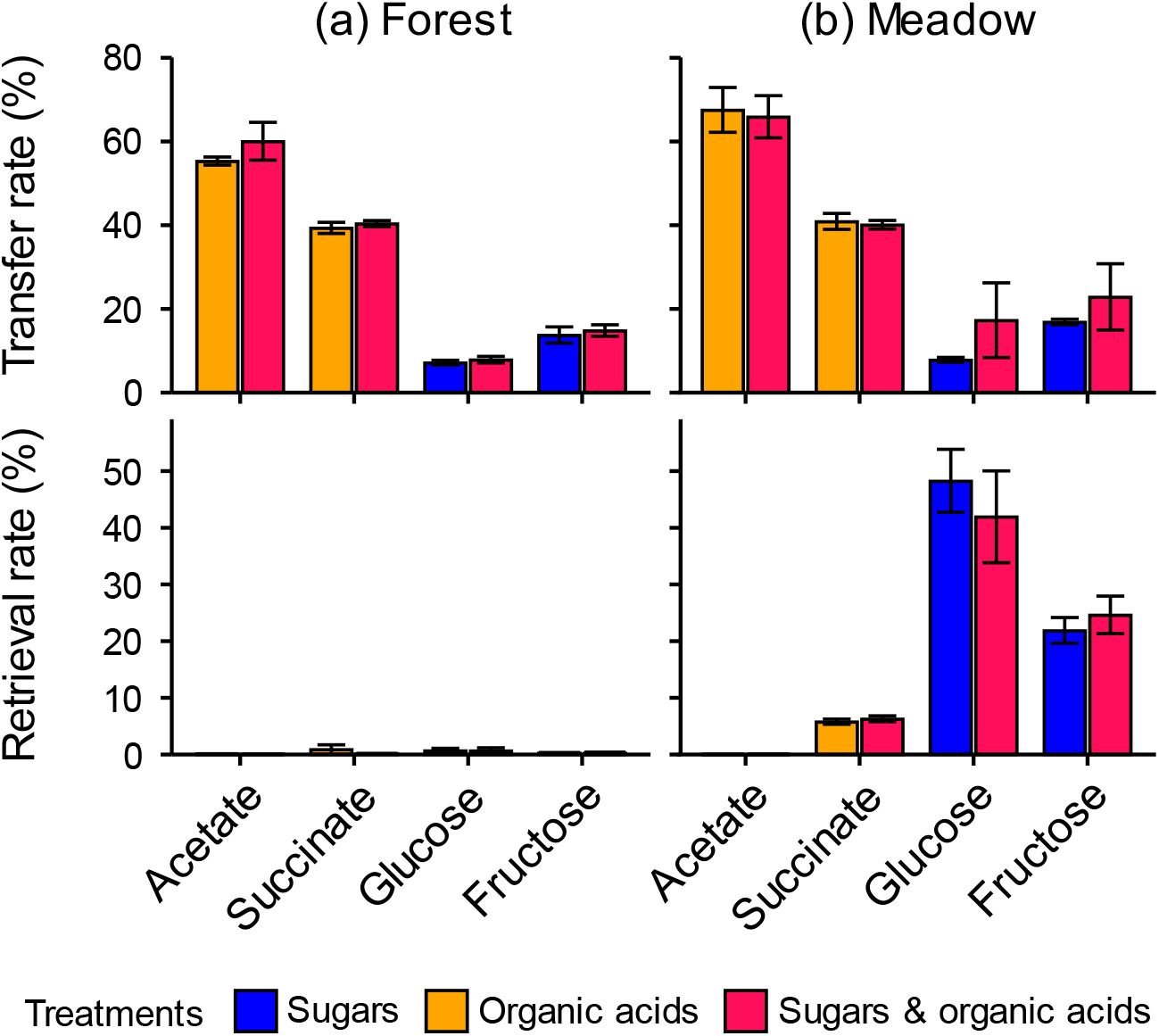
Transfer and retrieval rates (%) of substrates during and after a 6-hour artificial root exudate pulse in (**a**) forest and (**b**) meadow soils. The experiment included three labile substrate treatments (‘exudates’): sugar-only (blue), containing glucose and fructose; organic acid-only (orange), containing acetate and succinate; and a mixed exudate (pink), containing all four compounds. Each compound was present at a concentration of 500 µmol C l^-1^ in the artificial root exudates. Transfer rates indicate the percentage of these compounds released during the 6-hour pulse, while retrieval rates indicate the percentage of initially transferred substrates recovered from the soil in the 12 hours following the pulse. Transfer and retrieval rates of individual compounds did not significantly differ between substrate treatments (Kruskal-Wallis test). Data are presented as means ± SE (*n* = 5).

**Figure 3.**
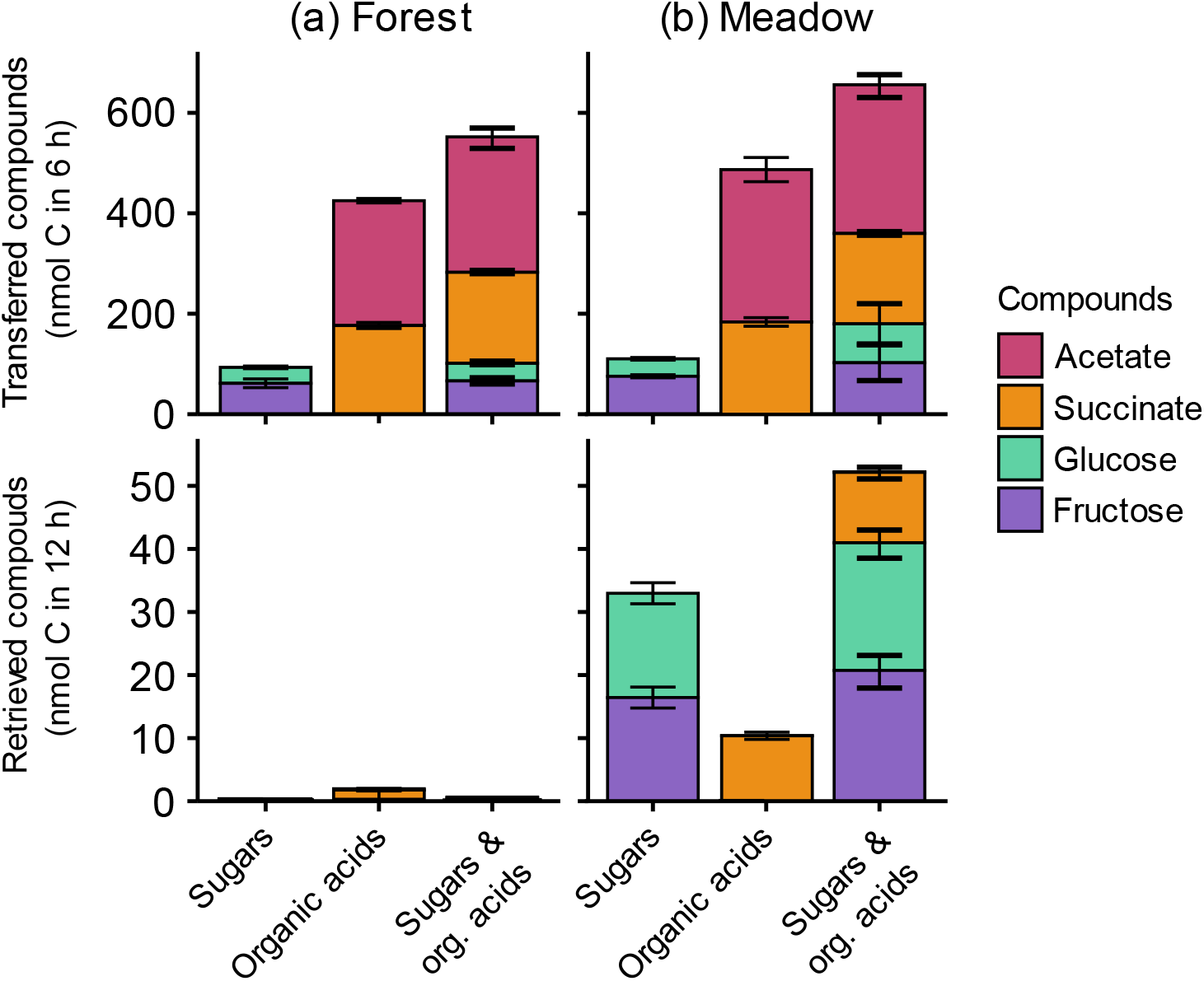
Substrate release and retrieval (nmol C) during and after a 6-hour artificial root exudate pulse in (**a**) forest and (**b**) meadow soils. The experiment included three labile substrate treatments (‘exudates’): sugar-only, containing glucose and fructose; organic acid-only, containing acetate and succinate; and a mixture exudate, containing all four compounds. Transferred compounds refers to the amount of C released during the 6-hour pulse, while retrieved compounds quantify the amount of initially added C recovered from the soil in the 12 hours following the pulse. Data are presented as means ± SE (*n* = 5).

In the hours after the labile substrate pulse had ended, less than 1% of the released compounds were back retrieved from the forest soil (Fig. 2). In the meadow soil, retrieval rates were varied for different compounds, i.e., less than 1% of acetate and about 6% of succinate were retrieved, while the retrieval rates of glucose and fructose were higher, at 28- 42% and 21-25% respectively (Fig. 2).

### 3.3 Organic acids, but not sugars, induced increased microbial metabolites and cations

The release of organic acids by reverse microdialysis (with or without sugars) led to a significant rise in the concentrations of the short-chain fatty acids (SCFAs) butyrate, lactate, and propionate during the second half of the simulated root exudation pulse (i.e., at about 3 hours after the start of the pulse) (Fig. 4, Tab. S2). Curiously, in the forest soil the increase in lactate levels was much more pronounced in response to the organic acids-only compared to the mixed exudate, with concentrations increasing 10-fold (Fig.4). In addition, organic acid release (with/ without sugars) significantly increased ammonium (NH4), potassium (K), magnesium (Mg), and calcium (Ca) concentrations around the probe during the pulse (Fig. 5, Tab. S2). The release of sugars, on the contrary, did not elicit any discernible changes in the concentrations of SCFAs or cations compared to the control (Fig. 4, Fig. 5, Tab. S2).

**Figure 4.**
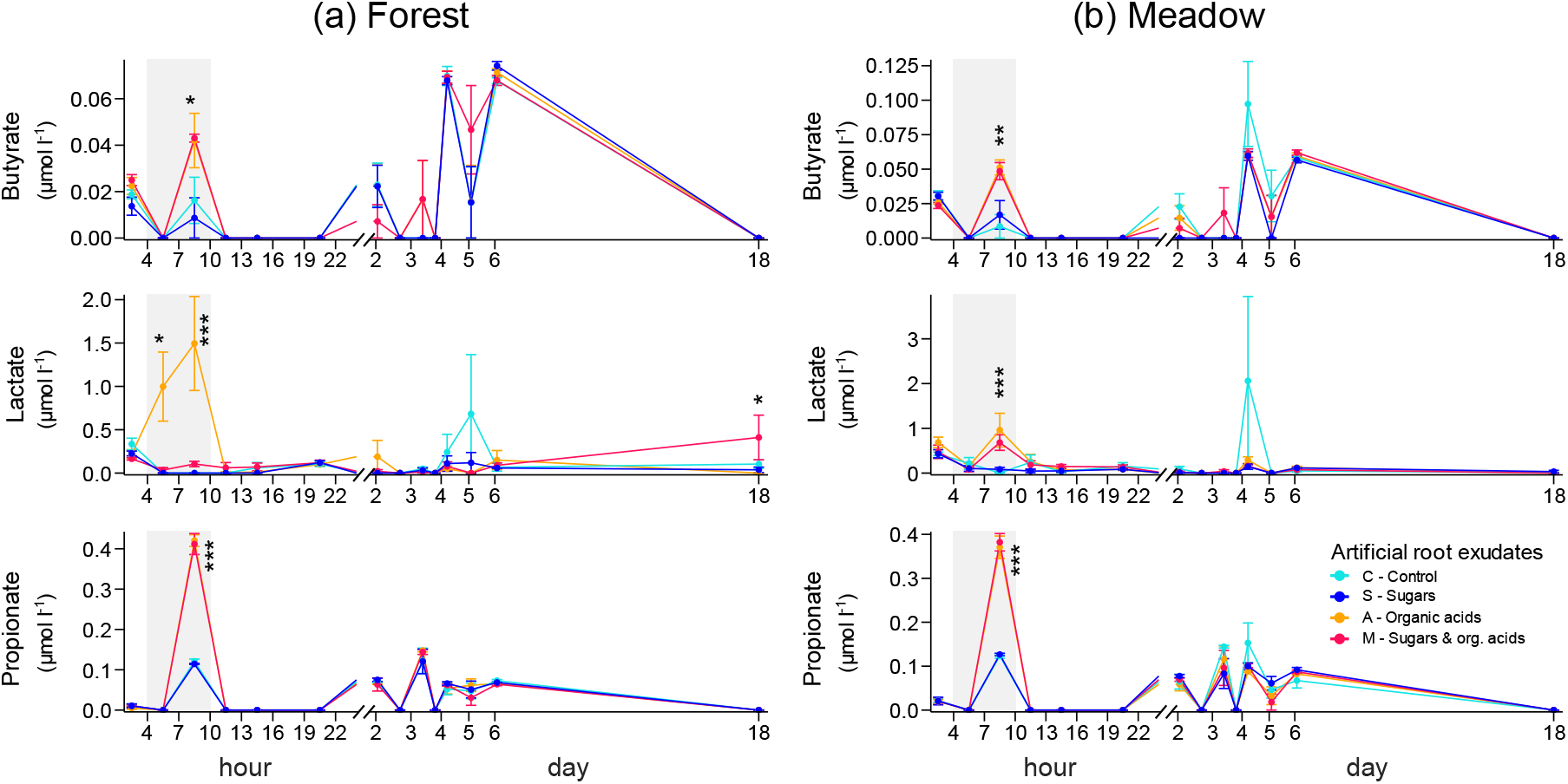
Concentration of butyrate, lactate, and propionate (µmol l^-1^) in dialysates collected from (**a**) forest and (**b**) meadow soils before, during and after a 6-hour long labile substrate pulse. The period of simulated exudation (hours 4-9) is indicated by a grey background. An x-axis break follows the initial 24-hour period, and subsequent observations (days 2-18) are presented on a day-timescale. Asterisks denote significant differences (Kruskal-Wallis test, *< 0.05, ** < 0.01, *** < 0.001) between soils that received an input of sugars (dark blue), organic acids (orange), a mixture of sugars and organic acids (pink), and control (light blue). Post-hoc test results (Dunn’s test) are provided in Table S3. Data are presented as means ± SE (*n* = 5). Each data point represents the mean concentration measured in dialysates collected over a 3-hour period on day 1 and a 4.5-hour period on subsequent days, plotted at the midpoint of each collection period.

**Figure 5.**
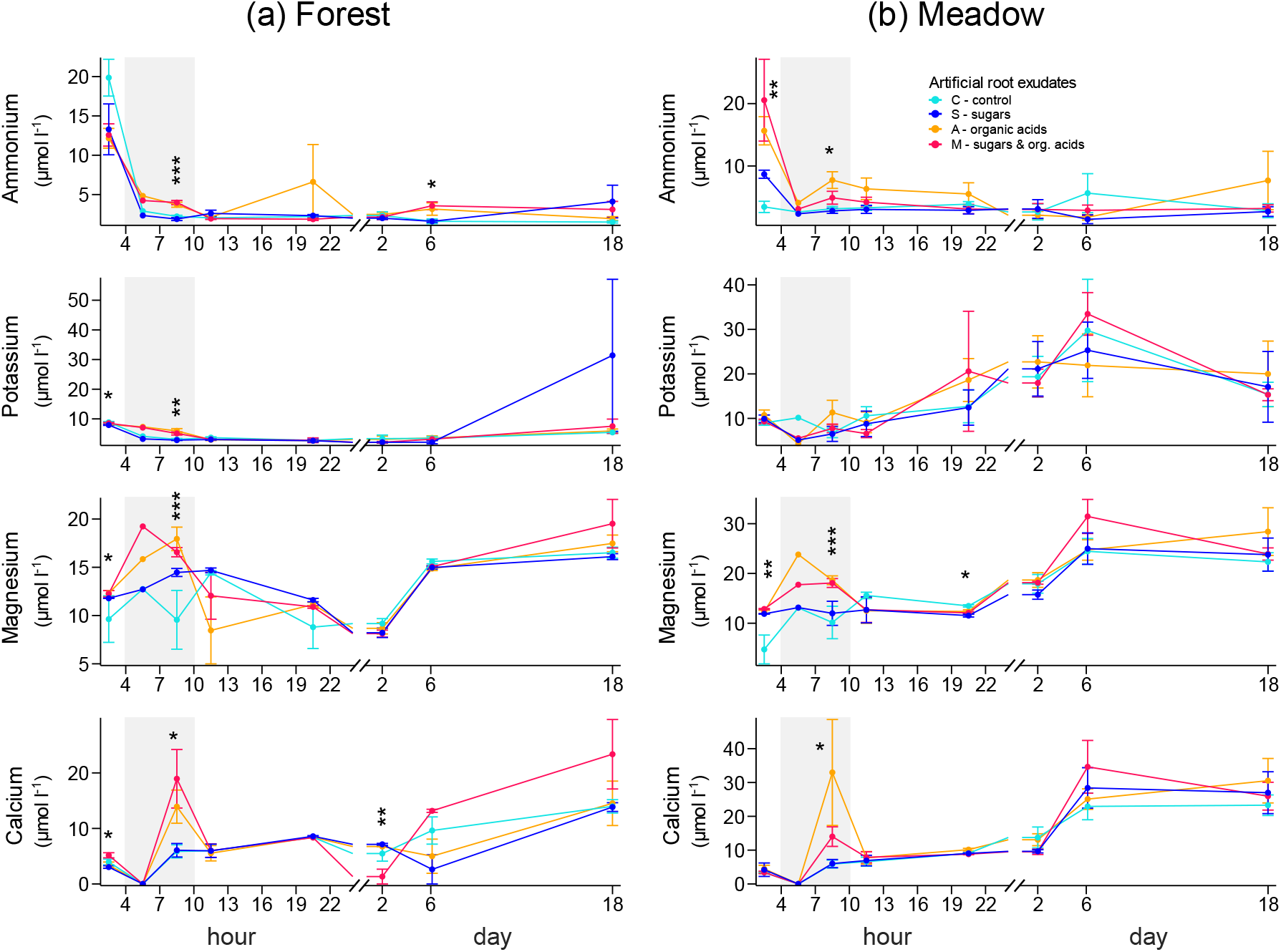
Concentrations of ammonium, potassium, magnesium, and calcium (µmol l^-1^) in dialysates collected from (**a**) forest and (**b**) meadow soils before, during and after a 6-hour long labile substrate pulse. The period of simulated exudation (hours 4-9) is indicated by a grey background. An x-axis break follows the initial 24-hour period, and subsequent observations (days 2-18) are presented on a day-timescale. Asterisks denote significant differences (Kruskal-Wallis test, * < 0.05, ** < 0.01, *** < 0.001) between soils that received an input of sugars (dark blue), organic acids (orange), a mixture of sugars and organic acids (pink), and control (light blue). Post-hoc test results (Dunn’s test) are provided in Table S3. Data are presented as means ± SE (*n* = 5), except for hour 4 (*n* = 1). Each data point represents the mean concentration measured in dialysates collected over a 3-hour period on day 1 and a 4.5-hour period on subsequent day, plotted at the midpoint of each collection period.

Providing sugars alongside organic acids did not affect the concentrations of metabolites or cations compared to organic acids-only treatment, except for the higher lactate levels in the latter (Fig.4). This pattern of organic acids inducing higher concentrations of metabolites and cations, while sugars did not, was largely consistent across both investigated soils.

In addition to the immediate effects following the simulated root exudation pulse, treatment-induced changes in metabolite and cation concentrations were also observed at later time points (day 2-18, Fig. 4, Fig. 5). They were, however, not consistent across soils or treatments. For instance, lactate concentration was increased after 18 days in the forest soil after mixed compound addition. Similarly, NH4 was significantly increased in forest soils 6 days after having received organic acids. There was, however, no effect on NH4 concentrations beyond the first 24 hours in the meadow treatment.

### 3.4 Combined sugars and organic acids are incorporated into microbial biomass; sugars-only into microbial storage

The sum of all microbial PLFAs (nmol C g^-1^ DW), which we used as a proxy for total microbial biomass, remained unaffected by simulated root exudation (>2.5 mm distance from membrane) across treatments (Fig. 6). Similarly, there was no increase in PLFA concentrations associated with any individual microbial group (gram-negative and gram-positive bacteria, fungi, actinobacteria) (Fig. S4). The sum of microbial NLFAs, which we use as a proxy for microbial storage compounds, significantly increased in meadow soils that had received a mix of organic acids and sugars (Fig. 6). This increase was primarily attributed to an increase in the general NLFA biomarkers, such as palmitic acid (16:0) (Fig. S6).

**Figure 6.**
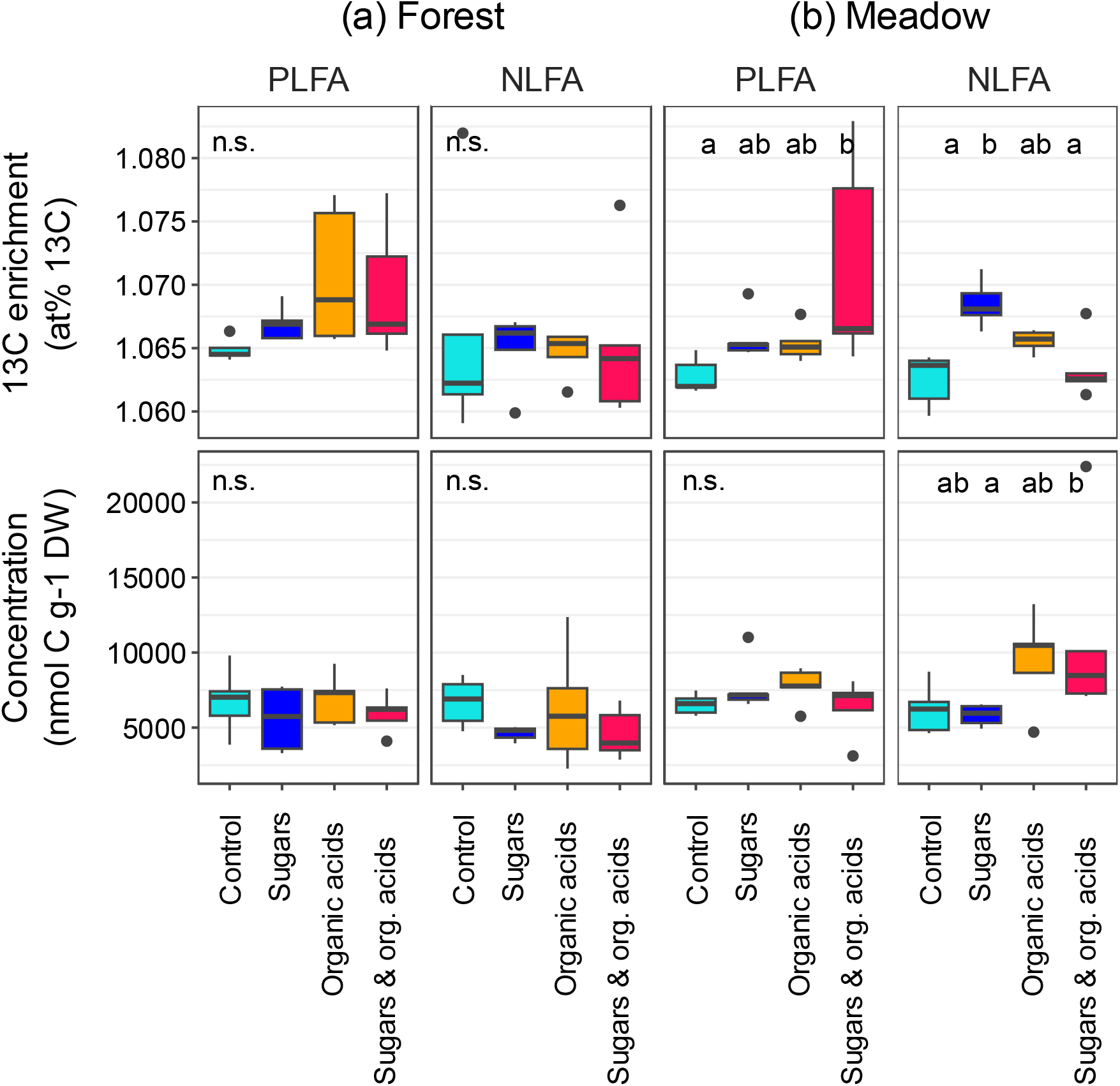
The concentration (nmol C g^-1^ dw) and ^13^C enrichment (at% ^13^C) of phospholipid fatty acids (PLFAs) and neutral lipid fatty acids (NLFA) in (**a**) forest and (**b**) meadow soils. Letters indicate significant difference between treatments (Kruskal-Wallis test, p < 0.05, Post-hoc test: Dunn’s test) that received only sugars (dark blue), only organic acids (orange), a mixture of sugars and organic acids (pink), and a control (light blue) that did not receive a labile substrate pulse (*n* = 5). Note that our sampling excluded soil within a 2.5 mm radius of the microdialysis membrane, making the measurements a conservative estimate of lipid-derived fatty acids enrichment in the immediate vicinity of the membrane.

PLFAs were significantly enriched in ^13^C in meadow soils in response to mixed labile substrate addition (at% ^13^C: Fig. 6). A similar, though statistically non-significant, trend of ^13^C incorporation in PLFAs was observed in forest soils following organic acid additions (with or without sugars). However, this trend was significant in forest soils when looking only at general and gram-negative PLFA biomarkers (Fig. S5). We observed a significant ^13^C enrichment into NLFAs after sugar addition in meadow soils (Fig. 6). Again, this enrichment was mainly driven by incorporation of ^13^C into general NLFA biomarkers, such as 16:0 (Fig S7).

## 4. Discussion

Organic acids and sugars are two of the major compound classes exuded by plant roots. It has often been assumed that sugars act as a primary energy source for microbes, while organic acids play a key role in releasing nutrients from mineral and humic surfaces (Keiluweit et al., 2015). However, our findings extend these traditional views by demonstrating that organic acids in plant root exudates may also serve as an essential energy source for microbes, which are metabolised even faster and to a greater extent compared to sugars.

### 4.1 Disparity in sugar and organic acid release

Our results show considerably higher passive release rates of organic acids compared to sugars in both soils (Fig. 2, Fig. 3). The already low release rates of sugars dropped further in the second half of the pulse, indicating rapid saturation of sugars around the microdialysis membrane (Fig S2). Moreover, in the forest soil, lower sugar transfer rates were associated with high back retrieval rates after the substrate pulse ended (Fig. 2). We consistently observed this pattern across our previous studies involving various soil types (König et al., 2022; Wiesenbauer et al., 2024) suggesting that lower rates of passive sugar release into soil compared to organic acids might be a widespread phenomenon. Our previous research demonstrated that higher substrate diffusion through microdialysis into intact soil can only be attributed to higher rates of biotic or abiotic compound removal around the membrane, which supersedes effects of molecular size and charge (König et al., 2022). Higher removal rates of acetate and succinate may partly be attributed to their charged nature, which facilitates rapid adsorption to minerals (Jones et al., 2003). Nevertheless, the significantly higher respiration derived from organic acids suggests a more substantial microbial uptake and mineralization rate compared to sugars (Fig. 1), which may have contributed to the consistently high release of organic acids during the simulated root exudation (Fig. S2). Our findings imply a microbial preference for organic acids over sugars, challenging the established view that sugars (monosaccharides) are the preferred source of readily available substrates which are rapidly consumed (Jones and Murphy, 2007; Gunina and Kuzyakov, 2015).

### 4.2 Preferential utilization of organic acids over sugars

Why do microbial communities in both soils prefer to assimilate organic acids, even though catabolizing sugars typically yields more energy? The full oxidation of hexoses, such as glucose and fructose during aerobic respiration yields about 28-32 ATP (Romano and Conway, 1996; Pastor et al., 2019), while those of organic acids such as acetate only yields about 12 ATP.

Despite this, we found that microbes favoured acetate and succinate. These organic acids are often considered metabolic endproducts in bacteria due to their partial oxidation, yet they serve as vital sources of C and energy that offer distinct advantages. For instance, acetate can be metabolized through the formation of acetyl-coenzyme A (acetyl-CoA), employing a shorter pathway that bypasses the decarboxylation step of glucose to acetyl-CoA (S. Zhang et al., 2019; Hosmer et al., 2023). This acetyl-CoA then enters the TCA cycle and is linked to energy production through a series of reactions that efficiently generate ATP and other energy carriers. Under conditions of high substrate flux, shorter metabolic pathways result in higher rates of ATP production, albeit at a lower ATP yield per unit of metabolised substrate (Kreft et al., 2020).

The choice to metabolise organic acids over sugars suggests a strategic trade-off by microbes between maximizing rate of ATP production and thus biomass growth and optimizing ATP yield (Kreft and Bonhoeffer, 2005; Wortel et al., 2018; Kreft et al., 2020). In environments where rapid biomass accumulation is crucial and substrate is in excess, such as in the rhizosphere or the here investigated exudation hotspots, the fast but lower-yielding ATP production from organic acid catabolism may be more beneficial than pursuing the higher ATP yield from sugar metabolism.

In addition, sugars and organic acids fuel two major opposing pathways of the central C metabolism: glycolysis, which breaks down glucose to pyruvate to produce energy, and gluconeogenesis, which synthesises glucose from non-carbohydrate sources (Schink et al., 2022). It is thought that microbes can either utilize glucose for biomass synthesis and energy production, or metabolize organic acids, since both C-metabolisms run in opposite directions (Schink et al., 2022). However, it has also been shown that E.coli co-consumes acetate and sugars, with acetate entering the TCA cycle during growth on glucose (Enjalbert et al., 2017). Given their faster diffusion, an initially higher organic acid concentration in the mixed substrate treatment might have signalled the microbial cells to opt for gluconeogenesis as the primary pathway in their central C metabolism. Usually, a lag phase occurs during the switch between glycolytic (sugars) to gluconeogenic (organic acids) substrates (Basan et al., 2020). This, together with the fact that ^13^C respiration rates were lowest when only sugars were released, suggests that gluconeogenesis rather than glycolysis may have been the default pathway in the majority of soil microbes. This aligns with the typical soil metabolic landscape, as a significant fraction of SOM is microbial necromass (i.e., proteins, storage and membrane lipids), whose breakdown products are gluconeogenic rather than glycolytic substrates.

### 4.3 Microbial incorporation of substrate-C into biomass

Our data indicates that microbes took up released exudates. The highest ^13^C enrichment was found in the combined sugar and organic acid treatment in the meadow soil, which could be attributed to the higher release of substrate-C in mixed exudate soils (Fig. 3). Additionally, in forest soils, general and gram-negative biomarkers showed significant ^13^C incorporation into PLFAs in the organic acid treatment, while sugars lead to a significant enrichment in NLFA biomarkers in the meadow soil. Overall, we observed only a relatively low ^13^C signal in PLFAs and NLFAs, which we attribute to our limitation of analysing soil only from beyond a >2.5 mm radius of the microdialysis membrane. Sampling soil from as close as possible to the membrane would yield clearer and more significant ^13^C enrichment pattern, as we have demonstrated before (Wiesenbauer et al., 2024).

### 4.4 Organic acids input causes short-chain fatty acids production

The assimilated substrates were directed towards different catabolic and anabolic processes, including the production of metabolites. Organic acids, with or without sugars, promoted microbial activity (as evident from ^13^C respiration measurements) and the production of short-chain fatty acids (SCFAs) such as butyrate, lactate, and propionate (Fig. 4). However, sugars alone did not induce metabolite production.

It remains unclear whether the butyrate, propionate and lactate we observed in the organic acid treatments originated from the breakdown of these organic acids (succinate) or from the induced breakdown of soil organic matter. SCFAs are commonly produced in anoxic environments as intermediates of organic matter degradation through fermentation. However, they can also occur as byproducts of high substrate turnover in aerobic conditions (Wolfe, 2005). Notably, SCFAs themselves are a non-fermentable C source and require a terminal electron-acceptor, such as O2 in aerobic or NO3 in anaerobic conditions, to be fully oxidized (Pavoncello et al., 2022). The anoxic sites conducive to fermentation may emerge at spots of high microbial activity, like (artificial) root exudation hotspots, where O2 consumption due to substrate degradation exceeds the O2 influx (Kreft et al., 2020). Indeed, it has been shown that O2 concentrations decreases during times of active root exudations in the rhizosphere of *Vicia Faba* (Garcia Arredondo et al., 2023) and after the release of glucose, acetic acid, and oxalic acid near an artificial root (Keiluweit et al., 2015). Under these O2- limited conditions, microbes perform shorter incomplete catabolic pathways such as fermentation, which support high growth rates due to higher ATP flux, despite their lower ATP yield compared to more yield-efficient pathways like aerobic respiration. This illustrates a trade-off between growth rate and yield (Kreft and Bonhoeffer, 2005; Wortel et al., 2018; Kreft et al., 2020).

Here, we hypothesise that an O2 depletion was caused primarily by the respiration of added substrate (Wiesenbauer et al., 2024), specifically organic acids. While SOM decomposition prompted by substrate addition might have contributed to O2 consumption, no significant changes in SOM-derived respiration were observed (Fig. 1). However, localised increases in SOM-derived respiration may have gone undetected.

Anaerobic respiration is increasingly recognised as a fundamental trait to colonise the rhizosphere (Lecomte et al., 2018). Until now, research on anaerobic metabolic processes has focussed on aqueous environments and paddy soils, leaving the rhizosphere of forest and meadow soil notably understudied (Lecomte et al., 2018). We observed butyrate, lactate, and propionate production in response to organic acid addition in our artificial rhizosphere (Fig. 4). While direct comparisons with rhizosphere studies are lacking, research on anaerobic metabolism in other environments provides context. For instance, in rice paddy soils, butyrate and propionate are common intermediary products of organic material decomposition (Rui et al., 2009). Similarly, in the gut, butyrate and propionate are commonly metabolised from complex carbohydrates like plant cell wall polysaccharides (Koh et al., 2016; Nogal et al., 2021), as well as from host-derived succinate by gut microbiota (Wei et al., 2023). In contrast to these environments, soils provide a wide array of alternative terminal electron acceptors in anaerobic conditions, such as NO3, SO2, or Fe (III), which would allow to feed intermediary fermentation products into anaerobic respiration (Keiluweit et al., 2017; Lecomte et al., 2018). One way to provide a more reliable evidence for anaerobic metabolism in the rhizosphere could be to monitor O2 levels simultaneously when simulating root exudation (Garcia Arredondo et al., 2023).

### 4.5 Organic acid induced mobilization of cations

The mobilization of K (only forest), Ca, and Mg concentrations in both soils (Fig. 5) could be a consequence of abiotic interactions between organic acids and soil minerals.

Organic acids can dissolve minerals through acidification, chelation, and exchange reactions (Adeleke et al., 2017), thereby making nutrients available for plant uptake (Giehl and von Wirén, 2014). Acidolysis occurs when protons from organic acids lower the pH, inducing the release of cations like Fe, K and Mg (Uroz et al., 2009; Adeleke et al., 2017). Moreover, organic acids form complexes with cations and mineral surfaces/metals in a process called chelation (Uroz et al., 2009), thereby increasing the solubility of nutrients like Ca, Mg, Fe, and Al (Golubev et al., 2006; Xu and Gao, 2008). Additionally, organic acids can displace cations through ligand exchange (Oburger et al., 2009; Keiluweit et al., 2015). Although acetate is a relatively weak complexing agent, it is possible that the protons it released mobilized cations such as K, Ca, Mg and NH4 from clay minerals or organic matter via cation exchange reactions.

As our findings demonstrate (Fig. 4), various bacteria and fungi produce and release organic acids similar to those produced by plant roots, which can fulfil the same function of solubilizing elements (Banfield et al., 1999; Jones et al., 2003; van Schöll et al., 2008; Adeleke et al., 2010; Uroz et al., 2011; R. A. Adeleke et al., 2012; R. Adeleke et al., 2012). Consequently, the observed increase in nutrient concentrations (K, Ca, Mg) in response to both added and microbially produced organic acids highlights their pivotal role in mobilizing mineral-associated nutrients in the rhizosphere.

### 4.6 Implications of rhizosphere feedback on root exudation

Root exudation of primary metabolites, such as sugars, organic acids and amino acids, is thought to occur through passive transport along electrochemical and concentration gradients between rhizodermal cells and the soil environment (Canarini et al., 2019). These gradients are affected by the rate at which compounds are removed by microbial activity or interaction with soil surfaces. Our findings suggest that while organic acids are quickly removed by both biotic and abiotic processes, sugars are not. However, if sugars are not readily metabolised by soil microbes, this impairs the plant’s ability to release photosynthates as sugars into the rhizosphere. Although it has been suggested that anions like organic acids tend to be released from roots more easily than sugars due to charge differences (Jones, 1998), our study is the first to demonstrate that soil microbes may contribute to this pattern by preferably taking up organic acids over sugars at root exudation hotspots.

Traditional root exudation measurements have reported sugars as a significant proportion of total root exudates (Farrar et al., 2003; Badri and Vivanco, 2009). However, these measurements typically involve removing surrounding soil and sampling root exudates hydroponically to avoid the alteration of exuded compounds by the surrounding soil matrix and microbes (Kuijken et al., 2015; Oburger and Schmidt, 2016). This approach, however, also eliminates soil microbial feedback on diffusional exudation rates, which may result in an overestimation of sugar exudation rates. Our research suggests that the exudation of sugars by roots may be impeded by low microbial metabolism of sugars in the rhizosphere, emphasising the importance of biotic and abiotic rhizosphere feedback mechanisms in regulating root exudation rates.

## Acknowledgements

We would like to thank Joshua Schimel for his careful reading and constructive feedback, which improved this paper.

## Data availability

Data to this article can be found online at https://doi.org/10.5281/zenodo.13338424.

## Declaration of generative AI and AI-assisted technologies in the writing process

During the preparation of this work the author(s) used ChatGPT in order to improve the readability and language of the manuscript. After using this tool, the author(s) reviewed and edited the content as needed and take full responsibility for the content of the published article.

