## Supplementary information for "Soil microbes prefer organic acids over sugars in simulated root exudation"

Article:

The following Supporting material is available for this article:

**Fig. S1** Photographs of experimental setup

**Fig. S2** Concentration of compounds released during the first and second half of the simulated exudation

**Fig. S3** Compound concentrations of perfusate compounds during day 1

**Fig. S4** Concentration of PLFAs for biomarker groups

**Fig. S5** $^{13}$C enrichment of PLFAs for biomarker groups

**Fig. S6** Concentration of NLFAs for biomarker groups

**Fig. S7** $^{13}$C enrichment of NLFAs for biomarker groups

**Table S1** Post-hoc test results of substrate-derived respiration

**Table S2** Post-hoc test results of compound concentrations

**Table S3** Post-hoc test results of perfusate compound concentrations

**Methods S1** Analysis of microbial biomass C and N

**Methods S2** High-performance liquid chromatography of sugars

**Methods S3** High-performance liquid chromatography of anions

**Methods S4** High-performance liquid chromatography of cations

**Methods S5** Calculation of respiration rates

**Methods S6** Phospholipid and neutral lipid extraction and analysis

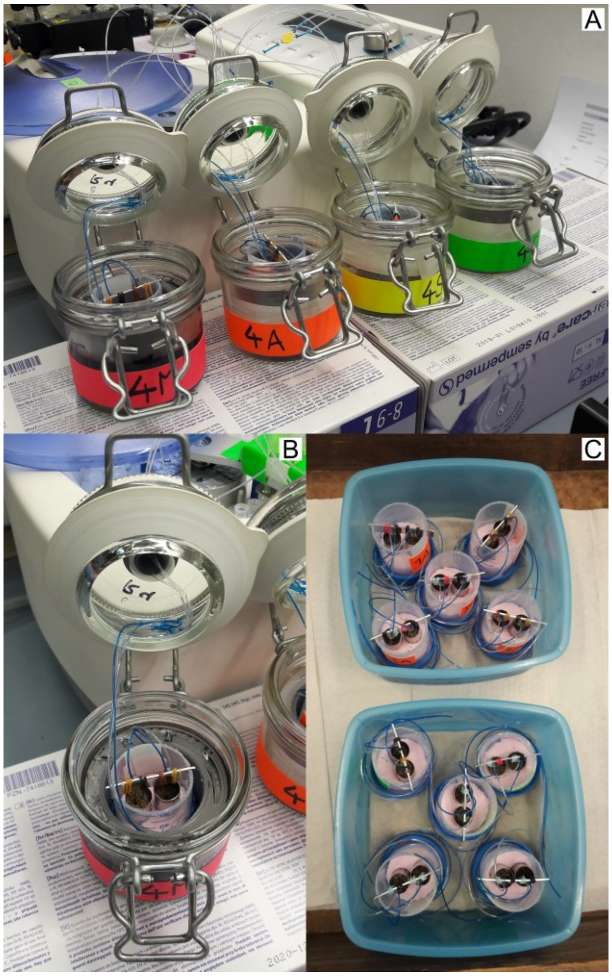


**Figure S1** These photographs illustrate our experimental setup. (**A**) Each mesocosm (cut of 50 ml falcon tube) held two small undisturbed soil cores (1 cm diameter, 3 cm height). The two microdialysis probes, one per soil core, were kept stable in the soil by attaching them to an aluminium strip on top of the falcon. The dialysate of the two soil cores per mesocosm were always pooled. (**B**) During microdialysis the mesocosms were kept in airtight jars fitted with a septum through which the microdialysis tubing was passed through. This allowed for simultaneous collection of dialysates in fraction collectors (in the background of picture A and B) and gas sampling with closed jar lid. (**C**) Between measurements the mesocosms were placed in plastic containers with lids, with a wet paper towel on the bottom to reduce drying at room temperature. They were aired at least once a day and the weight of the mesocosms was tracked and water added on top of cores to adjust for water loss.

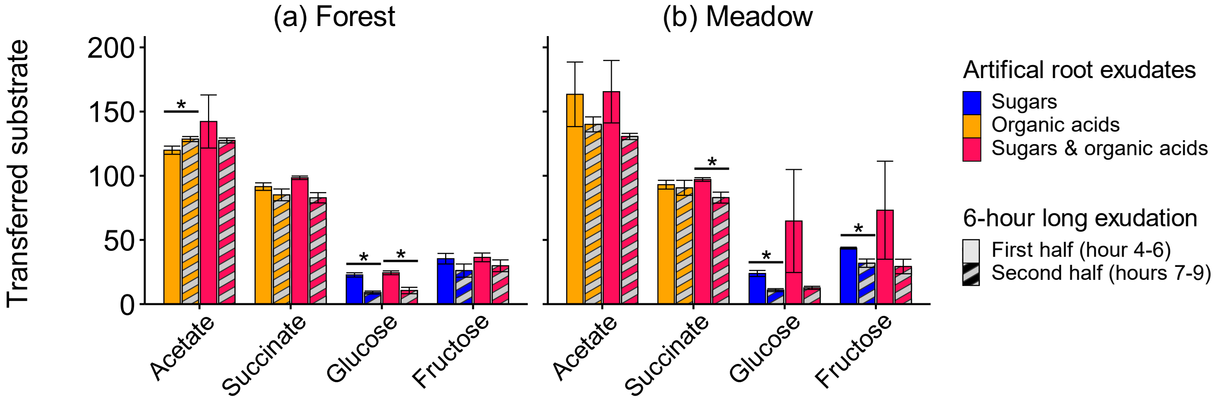


**Figure S2** The concentration (nmol C) of compounds released into (**a**) forest and (**b**) meadow soil, given for the first (solid colour, hours 4-6) and second half (dashed colour, hours 7-9) of the 6-hour long input pulse. Transferred compound concentrations are given for the artificial root exudates composed of either only sugars (blue), only organic acids (orange) or a mixture of sugars and organic acids (pink). All measures are means ± SE (*n* = 5). Asterisks indicate significant differences (Kruskal-Wallis test, $p < 0.05$) in released C between first and second half of input pulse.

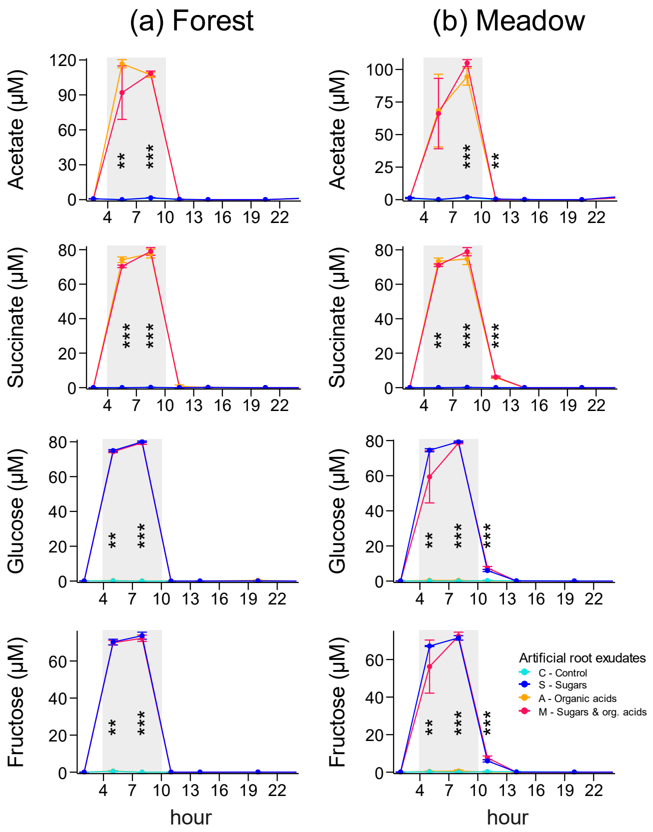


**Figure S3** Compound concentration (µmol $l^{-1}$) of organic acids (acetate, succinate) and sugars (glucose, fructose) that made up the perfusate ("artificial root exudate"). Concentrations were measured in dialysates collected from (**a**) forest and (**b**) meadow soils before, during and after a 6-hour long labile substrate pulse, highlighted with a grey background. It should be noted that the concentrations depicted during the pulse (orange background) are concentrations in dialysate. Which means that only the difference to the perfusate concentration (equivalent to 250 µM acetate, 125 µM succinate, 83.3 µM glucose, 83.3 µM fructose: each 500 µmol C $l^{-1}$) was transferred into soil. All measures are mean ± SE (*n* = 5). Asterisks indicate significant differences (Kruskal-Wallis test, * < 0.05, ** < 0.01, *** < 0.001) between soils that received an input of sugars (S: dark blue), organic acids (A: orange), a mixture of sugars and organic acids (M: pink) or control (C: green). Post hoc test results can be found in Table S3 (Dunn's test). Each data point represents the mean concentration measured in dialysates collected over a 3-hour period, with the point plotted at the midpoint of this period.

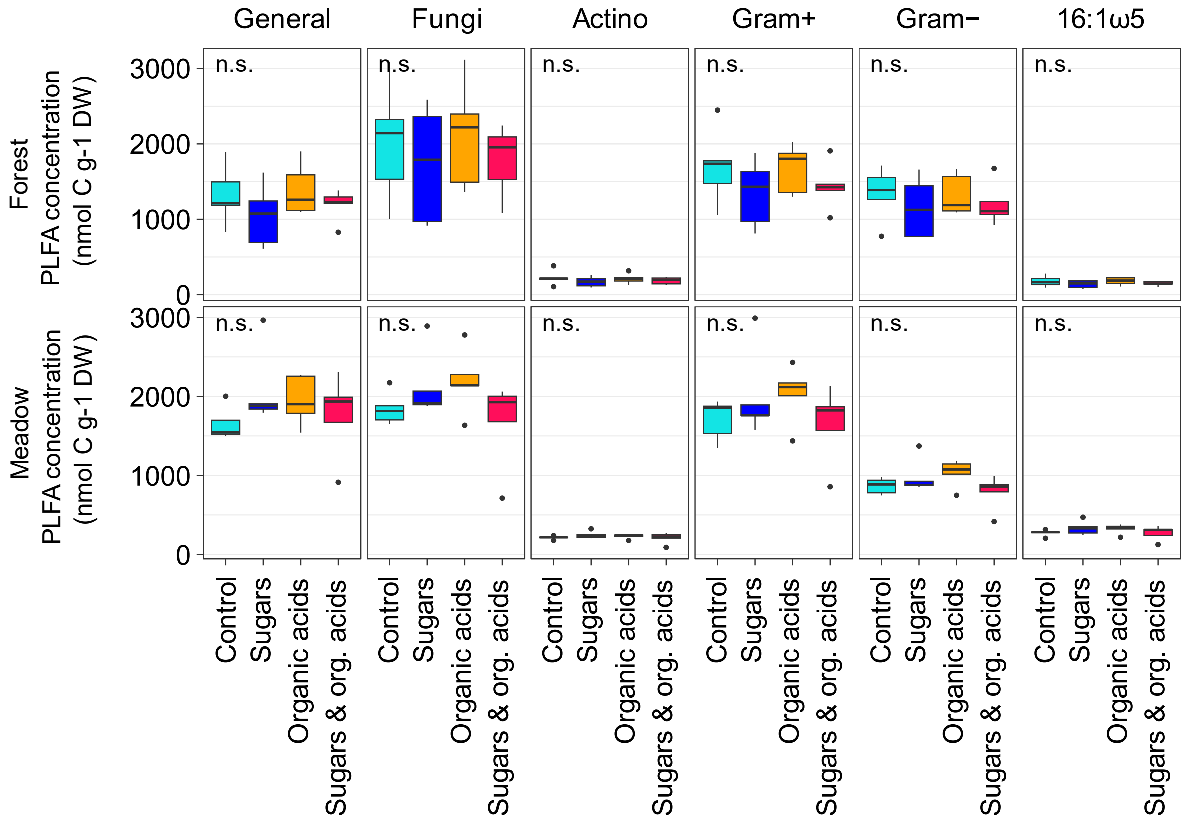


**Figure S4** The concentration (nmol C $g^{-1}$ dw) of phospholipid fatty acids (PLFAs) in forest and meadow soils. The fatty acids were grouped into general fatty acids, fungi, *Actinobacteria*, gram-positive bacteria (excluding *Actinobacteria*), gram-negative bacteria. The fatty acid 16:1ω5 was left ungrouped, because it is known to be a biomarker specific for arbuscular mycorrhiza fungi in meadow soil, but not in forest soil. Letters indicate significant difference between the treatments (Kruskal-Wallis test, $p < 0.05$, post-hoc test: Dunn's test) that received only sugars (dark blue), only organic acids (orange), a mixture of sugars and organic acids (pink) and a control (light blue) that did not receive a labile substrate pulse ($n$ = 5).

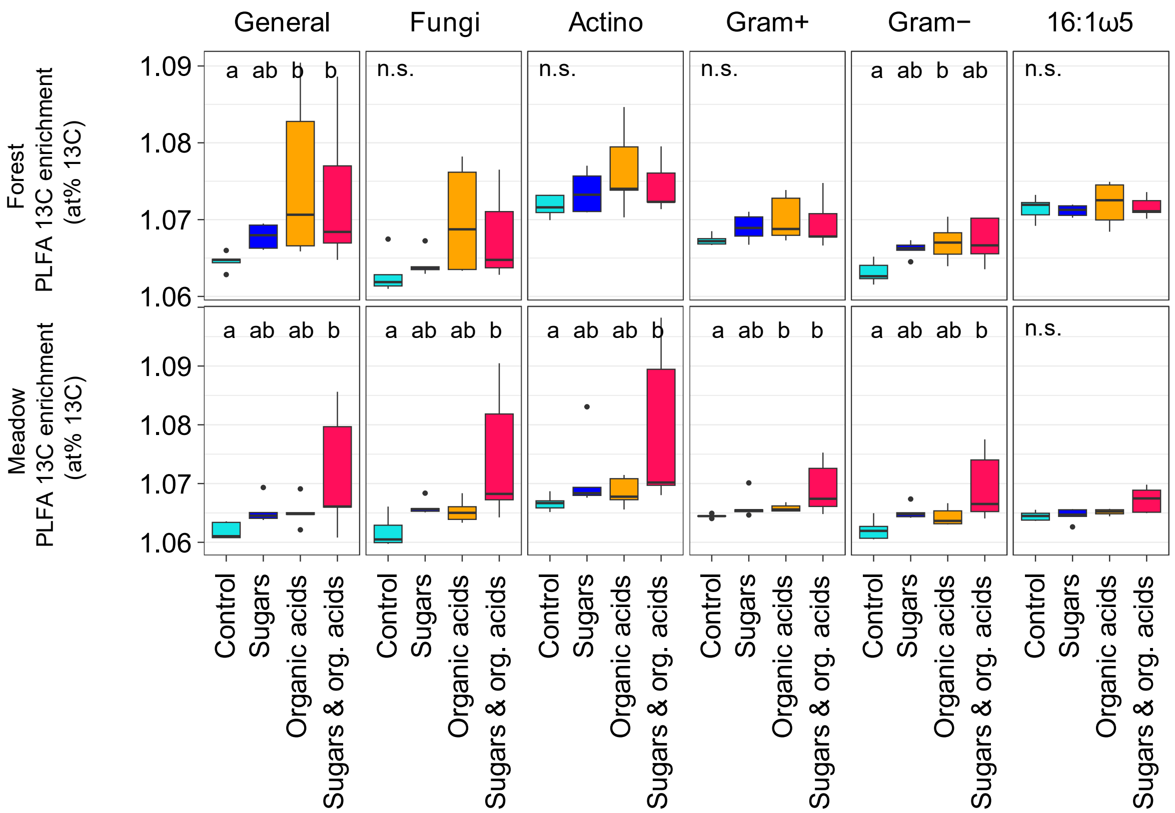


**Figure S5** The $^{13}C$ enrichment (at% $^{13}C$) of phospholipid fatty acids (PLFAs) in forest and meadow soils. The fatty acids were grouped into general fatty acids, fungi, *Actinobacteria*, gram-positive bacteria (excluding *Actinobacteria*), gram-negative bacteria. The fatty acid 16:1ω5 was left ungrouped, because it is known to be a biomarker specific for arbuscular mycorrhiza fungi in meadow soil, but not in forest soil. Letters indicate significant difference between the treatments (Kruskal-Wallis test, $p < 0.05$, post-hoc test: Dunn's test) that received only sugars (dark blue), only organic acids (orange), a mixture of sugars and organic acids (pink) and a control (light blue) that did not receive a labile substrate pulse ($n$ = 5).

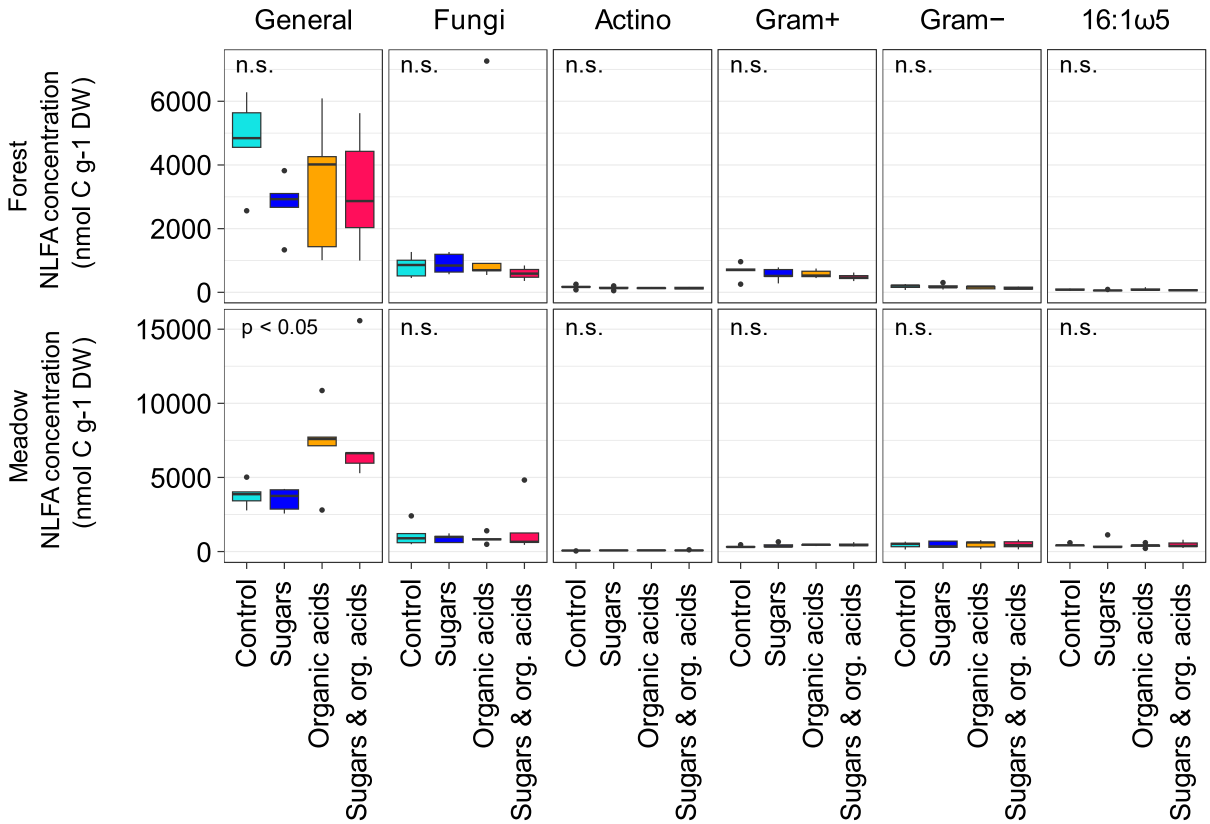


**Figure S6** The concentration (nmol C $g^{-1}$ dw) of neutral lipid fatty acids (NLFAs) in forest and meadow soils. The fatty acids were grouped into general fatty acids, fungi, *Actinobacteria*, gram-positive bacteria (excluding *Actinobacteria*), gram-negative bacteria. The fatty acid 16:1ω5 was left ungrouped, because it is known to be a biomarker specific for arbuscular mycorrhiza fungi in meadow soil, but not in forest soil. Letters indicate significant difference between the treatments (Kruskal-Wallis test, $p < 0.05$, post-hoc test: Dunn's test) that received only sugars (dark blue), only organic acids (orange), a mixture of sugars and organic acids (pink) and a control (light blue) that did not receive a labile substrate pulse ($n$ = 5).

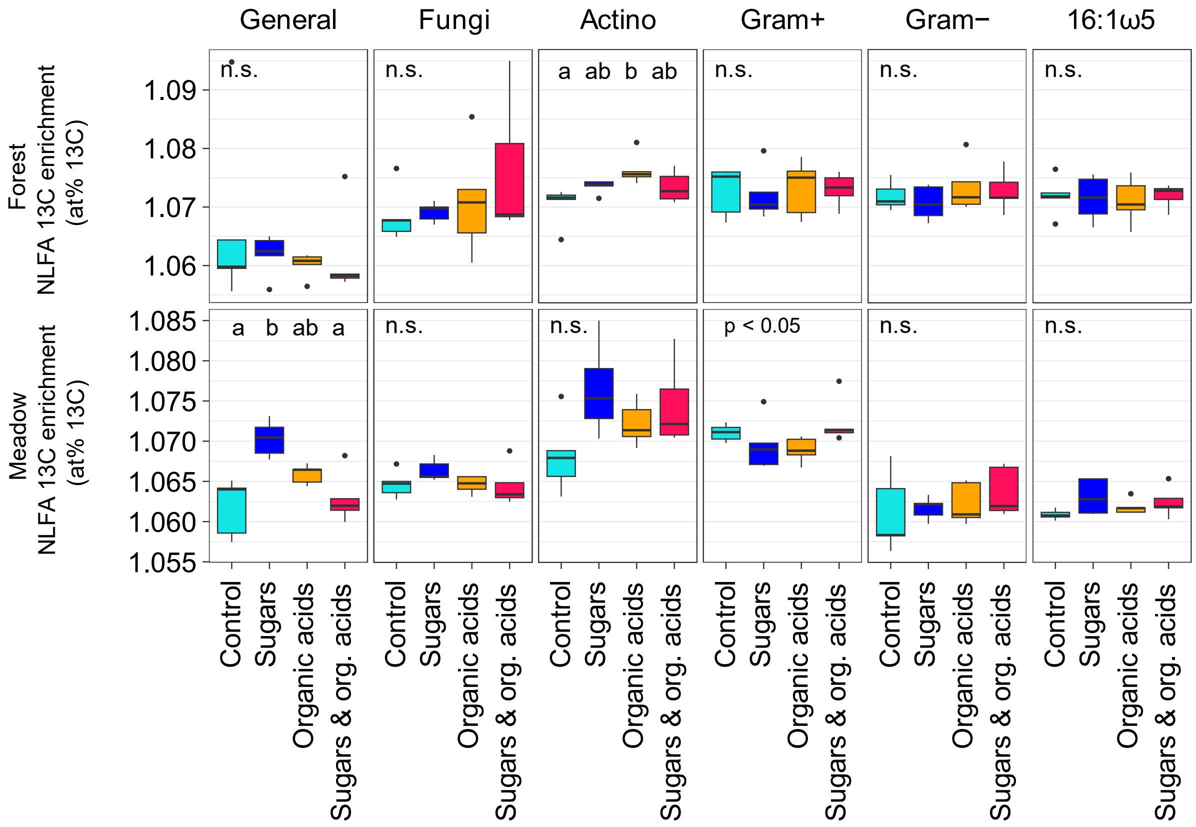


**Figure S7** The [13]C enrichment (at% [13]C) of neutral lipid fatty acids (NLFAs) in forest and meadow soils. The fatty acids were grouped into general fatty acids, fungi, *Actinobacteria*, gram-positive bacteria (excluding *Actinobacteria*), gram-negative bacteria. The fatty acid 16:1ω5 was left ungrouped, because it is known to be a biomarker specific for arbuscular mycorrhiza fungi in meadow soil, but not in forest soil. Letters indicate significant difference between the treatments (Kruskal-Wallis test, $p < 0.05$, post-hoc test: Dunn's test) that received only sugars (dark blue), only organic acids (orange), a mixture of sugars and organic acids (pink) and a control (light blue) that did not receive a labile substrate pulse ($n$ = 5).

**Table S1** Post-hoc test results (Dunn's test ) of substrate-derived respiration (Fig. 1).

| | (a) Forest | | | (b) Meadow |
|---|---|---|---|---|
| Day | 1 | | 2 | 1 |
| Hour | 6.5 | 11 | 26 | 6.5 |
| Substrate-respiration | $S^a$ $A^b$ $M^b$ | $S^a$ $A^b$ $M^b$ | $S^a$ $A^{ab}$ $M^b$ | $S^a$ $A^{ab}$ $M^b$ |

Superscript letters indicate significant differences in substrate-derived respiration between treatments that had received sugars (**S**), organic acids (**A**), and sugars combined with organic acids (**M**) as labile substrate pulse in (**a**) forest and (**b**) meadow soils.

**Table S2** Post-hoc test results (Dunn's test) of compound concentrations (Fig. 4, Fig. 5).

| | (a) Forest | | | | | | (b) Meadow | | | |
|---|---|---|---|---|---|---|---|---|---|---|
| Day | 1 | | | 3 | 6 | 18 | 1 | | | 6 |
| Hour | 1 | 4-6 | 7-9 | | | | 1 | 7 | 19 | |
| Butyrate | - | | $C^{ab} S^{a} A^{b} M^{ab}$ | | $C^{a} S^{b} A^{ab} M^{a}$ | | - | $C^{a} S^{a} A^{b} M^{ab}$ | | - |
| Lactate | - | $C^{a} S^{a} A^{b} M^{ab}$ | $C^{a} S^{a} A^{b} M^{ab}$ | | - | $C^{a} S^{b} A^{b} M^{a}$ | - | $C^{a} S^{a} A^{b} M^{b}$ | - | $C^{a} S^{ab} A^{b} M^{ab}$ |
| Propionate | - | | $C^{a} S^{a} A^{b} M^{b}$ | | $C^{a} S^{ab} A^{ab} M^{b}$ | | - | $C^{a} S^{a} A^{b} M^{b}$ | | - |
| $NH_4^+$ | - | - | $C^{ac} S^{a} A^{bc} M^{b}$ | - | $C^{a} S^{a} A^{b} M^{b}$ | $C^{a} S^{b} A^{ab} M^{b}$ | $C^{a} S^{ab} A^{b} M^{b}$ | $C^{a} S^{a} A^{b} M^{ab}$ | - | - |
| K | $C^{a} S^{b} A^{ab} M^{ab}$ | - | $C^{a} S^{a} A^{b} M^{b}$ | - | - | - | - | - | - | - |
| Mg | $C^{ab} S^{a} A^{b} M^{ab}$ | - | $C^{a} S^{a} A^{b} M^{b}$ | - | $C^{a} S^{b} A^{b} M^{ab}$ | - | $C^{a} S^{ab} A^{ab} M^{b}$ | $C^{a} S^{a} A^{b} M^{b}$ | $C^{a} S^{b} A^{ab} M^{ab}$ | - |
| Ca | $C^{ab} S^{a} A^{ab} M^{b}$ | | $C^{a} S^{ab} A^{ab} M^{b}$ | $C^{a} S^{a} A^{a} M^{b}$ | $C^{ab} S^{a} A^{a} M^{b}$ | - | - | $C^{a} S^{ab} A^{b} M^{ab}$ | - | - |

Superscript letters indicate significant differences between control (**C**) and treatments that had received sugars (**S**), organic acids (**A**), and sugars combined with organic acids (**M**) as labile substrate pulse in (**a**) forest and (**b**) meadow soils.

**Table S3** Post-hoc test results (Dunn's test) of concentrations of compounds present in the perfusate (Fig. S3).

| | (a) Forest | | (b) Meadow | | |
|---|---|---|---|---|---|
| Day | 1 | | 1 | | |
| Hour | 4 | 7 | 4 | 7 | 10 |
| Acetate | $C^{a}$ $S^{a}$ $A^{b}$ $M^{ab}$ | $C^{a}$ $S^{a}$ $A^{b}$ $M^{b}$ | - | $C^{a}$ $S^{ac}$ $A^{bc}$ $M^{b}$ | $C^{ab}$ $S^{a}$ $A^{b}$ $M^{b}$ |
| Succinate | $C^{a}$ $S^{ac}$ $A^{b}$ $M^{bc}$ | $C^{a}$ $S^{a}$ $A^{b}$ $M^{b}$ | $C^{a}$ $S^{ac}$ $A^{b}$ $M^{bc}$ | $C^{a}$ $S^{ac}$ $A^{bc}$ $M^{b}$ | $C^{a}$ $S^{a}$ $A^{b}$ $M^{b}$ |
| Glucose | $C^{a}$ $S^{b}$ $A^{ac}$ $M^{bc}$ | $C^{a}$ $S^{b}$ $A^{ac}$ $M^{bc}$ | $C^{a}$ $S^{b}$ $A^{ab}$ $M^{b}$ | $C^{a}$ $S^{b}$ $A^{ac}$ $M^{bc}$ | $C^{a}$ $S^{b}$ $A^{a}$ $M^{b}$ |
| Fructose | $C^{a}$ $S^{b}$ $A^{a}$ $M^{b}$ | $C^{a}$ $S^{b}$ $A^{a}$ $M^{b}$ | $C^{a}$ $S^{b}$ $A^{ab}$ $M^{b}$ | $C^{a}$ $S^{b}$ $A^{a}$ $M^{b}$ | $C^{a}$ $S^{b}$ $A^{a}$ $M^{b}$ |

Superscript letters indicate significant differences between control (**C**) and treatments that had received sugars (**S**), organic acids (**A**), and sugars combined with organic acids (**M**) as labile substrate pulse in (**a**) forest and (**b**) meadow soils.

*Post-hoc test was not able to detect differences.

**Methods S1 Analysis of microbial biomass C and N**

We analysed microbial biomass C by chloroform fumigation-extractions (Vance et al., 1987). Soils (2 g) were fumigated over chloroform for 24 h, and subsequently fumigated and non-fumigated soils were extracted with 15 ml KCl (1 M) for 30 min before filtering through ash-free cellulose filters (Whatman). KCl-extracts were measured with TOC/TN analyser (TOC-VCPH/TMN-1, Shimadzu, Japan), and microbial biomass C and N were calculated as the difference in dissolved organic carbon (DOC) and total nitrogen (TN) between fumigated non-fumigated soils. Two outlier values were removed from forest microbial biomass carbon. We used a conversion factor for microbial biomass C of 0.45 to take into account incomplete extraction (Joergensen, 1996). In forest soils, two outliers had to be removed from the microbial biomass C and C:N ratio.

**Methods S2 High-performance liquid chromatography to measure sugars**

We measured sugars in the dialysates using HPLC (Dionex ICS 5000+, Thermo Fisher, Germany). Sugars (glucose, fructose, galactose, sucrose) were measured on a Thermo CarboPac PA20 (0.4 x 150 mm) column with a Thermo CarboPac PA20G (0.4 x 35 mm) guard column at a constant flow rate of 8 µl $min^{-1}$ with a KOH solvent. The run was 25 min long, starting at 8 mM for 12 min, then increasing to 200 mM over 5 min. We kept the concentration at 200 mM for 1 minute, then lowered the concentration to 8 mM over 1 minute, keeping the concentration at 8 mM for 5 min until the end of the 25-minute run. While the analytical setup for sugar measurements was capable of detecting fructose, galactose, glucose, and sucrose, only glucose and fructose were present in our samples.

**Methods S3 High-performance liquid chromatography to measure anions**

We measured organic and inorganic anions in the dialysates using HPLC (Dionex ICS 5000+, Thermo Fisher, Germany). We analysed acetate, butyrate, citrate, formate, lactate, malate, nitrate, oxalate, phosphate, propionate, succinate, and sulfate. The anions were measured on a Dionex IonPac AS11-HC (2 x 250 mm) column with a Dionex IonPac AG11-HC (2 x 50 mm) guard column at a constant flow rate of 0.25 ml $min^{-1}$ with a KOH solvent. The 45 min HPLC run started at 1 mM for 10 min, then increased to 15 mM for 14 min (until min 24), then increased to 60 mM for 9 min (until min 33), where we kept it for 5 min at 60 mM (until min 38). Afterwards we lowered concentrations to 1 mM and kept it there until the end of the 45-minute run.

While the analytical setup for anion measurement was capable of detecting acetate, butyrate, formate, lactate, malate, oxalate, propionate, succinate, citrate, nitrate, phosphate, and sulfate, only butyrate, lactate, and propionate concentrations showed significant responses to the simulated exudation and are presented in the paper.

**Methods S4 High-performance liquid chromatography to measure cations**

We measured cations in the dialysates using HPLC (Dionex ICS 5000+, Thermo Fisher, Germany). We analysed ammonium, potassium, magnesium, manganese, and calcium. The cations were measured on a Dionex IonPac CS16 (5 x 250 mm) column with a guard column at a constant flow rate of 1 ml $min^{-1}$ with methanesulfonic acid as solvent. The dialysates from the first day were run with a short method and the remaining dialysates were measured with a longer method.

The shorter method was 35 minutes long, starting with 30 mM for 10 min, then increased to 60 mM over 5 min (until minute 15) at which concentration (60 mM) it remained for 10 min (until minute 25). Afterwards we decreased to 30 mM over 1 min (until min 26), at which concentration it remained until the end of the run at minute 35.

The longer method was 47 minutes long, starting with 30 mM for 22 min, then increased to 60 mM over 5 min (until minute 27) at which concentration (60 mM) it remained for 10 min (until minute 37). Afterwards we decreased to 30 mM over 1 min (until min 38), at which concentration it remained until the end of the run at minute 47.

While the analytical setup for cation measurement detected ammonium, potassium, magnesium, calcium, and manganese, though manganese levels could not be reliably quantified.

**Methods S5** Calculations of soil respiration rates and SOM- and substrate-derived respiration rates

The following equations were adopted from the supplements from König *et al.* (2022).

We used the following equations to calculate respiration rates and its $^{13}C$ signature:

$$CO_{2\ corrected\ t0} = \frac{CO_{2\ sample} \times \left(V_{jar} - V_{sample}\right) + CO_{2\ blank} \times V_{jar}}{V_{jar}}$$

where $CO_{2\ corrected\ t0}$ is the $CO_2$ concentration (ppm) at time 0, $CO_{2\ sample}$ and $CO_{2\ blank}$ refers to the measured $CO_2$ concentration (ppm) in sample or blank, $V_{sample}$ and $V_{jar}$ refers to the volume of the sampled gas and the glass jar.

$$^{13}CO_{2\ corrected\ t0} = \frac{CO_{2\ sample} \times \left(\frac{at\%\ ^{13}C_{sample}}{100}\right) \times \left(V_{jar} - V_{sample}\right) + CO_{2\ blank} \times \left(\frac{at\%\ ^{13}C_{blank}}{100}\right) \times V_{sample}}{V_{jar}}$$

where $^{13}CO_{2\ corrected\ t0}$ is the calculated concentration of $^{13}CO_2$ concentration (ppm) present in the jar after taking out sample t0 and replacement by the artificial air; at% $^{13}C_{sample}$ and at% $^{13}C_{blank}$ refers to the relative $^{13}C$ content (in at% $^{13}C$) in the measured gas sample t0 and the artificial air (blank).

$$^{13}CO_{2\ sample\ t1} = CO_{2\ sample\ t1} \times \frac{at\%\ ^{13}C_{sample\ t1}}{100}$$

where $^{13}CO_{2\ sample\ t1}$ is the relative concentration of $^{13}C$-$CO_2$ (ppm) in sample t1, $CO_{2\ sample\ t1}$ is the concentration of $CO_2$ at t1 (1 hour after t0).

$$CO_{2\ increase} = CO_{2\ sample\ t1} - CO_{2\ corrected\ t0}$$

$$^{13}CO_{2\ increase} = {}^{13}CO_{2\ sample\ t1} - {}^{13}CO_{2\ sample\ t0}$$

where $CO_{2\ increase}$ and $^{13}CO_{2\ increase}$ stand for the increase in total $CO_2$ and $^{13}CO_2$ (ppm), respectively, during the incubation time of 1 hour.

$$Respiration_{sample} = \left(\frac{\frac{CO_{2\ increase}}{10^6} \times V_{jar}}{Mol.Volume\ CO_2} \times 1000\right) \Big/ gDW$$

where $Respiration_{sample}$ refers to the total respiration rate in the samples that received labelled substrates (treatment) or those which did not (control), expressed in nmol $CO_2$ $h^{-1}$ $g^{-1}$ dry soil. Mol.Volume $CO_2$ represents the molecular volume of $CO_2$ at normal atmospheric pressure and room temperature (24.465 × $10^{-3}$ ml $\mu mol^{-1}$).

$$at\%\ ^{13}C_{respiration} = \frac{^{13}CO_{2\ increase}}{CO_{2\ increase}} \times 100$$

At% $^{13}C_{respiration}$ represents the relative $^{13}C$ content in respired $CO_2$ after substrate addition (in at% $^{13}C$).

$$APE\ ^{13}C_{respiration} = at\%\ ^{13}C_{treatment} - at\%\ ^{13}C_{control}$$

where at% $^{13}C_{treatment}$ and at% $^{13}C_{control}$ refer to the at% $^{13}C$ of respired $CO_2$ (as calculated with the previous formula) in the sample which received $^{13}C$-labelled substrate (treatment) and the corresponding one which did not (control).

$$Substrate\ Respiration = \frac{APE\ {}^{13}C_{respiration}}{APE\ {}^{13}C_{substrate}} \times Respiration_{treatment}$$

where Substrate Respiration represents the $CO_2$ production from the added substrate in nmol $CO_2$ $h^{-1}$ $g^{-1}$ dry soil. APE $^{13}C$ substrate stands for the $^{13}C$ enrichment of used substrate.

$$SOM\ Respiration_{treatment} = Respiration_{treatment} - Substrate\ Respiration$$

where SOM Respiration represents soil organic matter (SOM) derived respiration in nmol $CO_2$ $h^{-1}$ $g^{-1}$ dry soil.

**Methods S6 Phospholipid and neutral lipid extraction and analysis**

The lipids in the lyophilized "outer" soils were extracted with a monophasic mixture of chloroform, methanol and citrate buffer (pH = 4) in a 1:2:0.8 volume ratio (Bligh and Dyer, 1959; Frostegård et al., 1991). After phase separation with addition of chloroform and citrate buffer, the lower phase containing the total lipids was collected. These were consequently fractionated with silica solid phase extraction columns ( Phenomenex, Strata SI-1 Silica, 55 µm, 70 A) by eluting consecutively with a chloroform-ethanol mix (v:v = 98:2; Gorka *et al.*, 2023) yielding neutral lipids, pure acetone, and a chloroform-methanol-water mix (v:v:v = 5:5:1; Buyer & Sasser, 2012) yielding phospholipids. Phospho- and neutral lipids were derivatized to fatty acid methyl esters by mild alkaline methanolysis. The resulting PLFAs and NLFAs were analysed on a gas chromatograph (Trace GC Ultra, Thermo Scientific, Germany) coupled to a mass spectrometer (ISQ, Thermo Scientific, Germany) for fatty acid identification and quantification, and on a GC-Ultra (Thermo Fisher Scientific, Milan, Italy) coupled to an isotope ratio mass spectrometer (IRMS; Finnigan Delta-V, Thermo Fisher Scientific, Bremen, Germany) via a GC IsoLink (Thermo Fisher Scientific, Bremen, Germany) for determination of isotopic $^{13}C/^{12}C$ ratios. We used nonadecanoic acid (FAME 19:0) as an internal standard for quantification, and bacterial and fungal fatty acid methyl esters (BAME CP mix, Supelco; 37 Component FAME mix, Supelco) as qualitative external standards. Fatty acids with ≥20 carbon atoms and unidentified fatty acids were removed from the data.

Fatty acids were classified based on their taxonomic specificity into general microbial biomass (10:0, 14:0, 15:0, 16:0, 17:0, 18:0), fungi (18:1ω9c, 18:1ω9t, 18:2ω6,9, 18:3ω6,9,12), *Actinobacteria* (10Me17:0, 10Me18:0), gram-positive bacteria (i14:0, a15:0, i15:0, 16:0Me, i16:0, 17:0Me, a17:0, cy17:0, i17:0, i18:0), and gram-negative bacteria (14:1ω5, 15:1ω5, 15:1ω5c, 2OH 16:0, i16:1ω6, 16:1ω7, 16:1ω9, 16:2ω6,9, 17:1ω7, 17:1ω8, 18:1ω5, 18:1ω7, cy19:0, 19:1ω9)(Willers et al., 2015). The fatty acid

16:1ω5 was analysed separately as it is found in high abundances in spores of arbuscular-mycorrhizal fungi (particularly NLFAs) which we expected to be abundant in the meadow soil, but can also be found in gram-negative bacteria (Lekberg et al., 2022; Olsson and Lekberg, 2022).
